# Reproducible Coactivation Patterns of Functional Brain Networks Reveal the Aberrant Dynamic State Transition in Schizophrenia

**DOI:** 10.1101/2021.03.29.437611

**Authors:** Hang Yang, Hong Zhang, Xin Di, Shuai Wang, Chun Meng, Lin Tian, Bharat Biswal

## Abstract

It is well documented that massive dynamic information is contained in the resting-state fMRI. Recent studies have identified recurring states dominated by similar coactivation patterns (CAP) and revealed their temporal dynamics. However, the reproducibility and generalizability of the CAP analysis is unclear. To address this question, the effects of methodological pipelines on CAP are comprehensively evaluated in this study, including preprocessing, network construction, cluster number and three independent cohorts. The CAP state dynamics are characterized by fraction of time, persistence, counts, and transition probability. Results demonstrate six reliable CAP states and their dynamic characteristics are also reproducible. The state transition probability is found to be positively associated with the spatial similarity. Furthermore, the aberrant CAP in schizophrenia has been investigated by using the reproducible method on three cohorts. Schizophrenia patients spend less time in CAP states that involve the fronto-parietal network, but more time in CAP states that involve the default mode and salience network. The aberrant dynamic characteristics of CAP are correlated with the symptom severity. These results reveal the reproducibility and generalizability of the CAP analysis, which can provide novel insights into the neuropathological mechanism associated with aberrant brain network dynamics of schizophrenia.

**Highlights:**

1. Three coactivation patterns (CAPs) pairs with opposite coactivation profiles were identified, and the between-state transition probability was positively correlated with their spatial similarity.
2. Good spatial and temporal reproducibility and generalizability of CAPs were achieved under varied analytic methods and independent cohorts.
3. Schizophrenia patients showed altered temporal dynamics not only within the triple-network but also other primary and higher-order networks.

## 1. Introduction

Since the resting-state fMRI was confirmed to be physiologically meaningful,(B. Biswal et al., 1995) a number of resting-state fMRI studies have emerged, of which functional connectivity (FC) is one popular method to detect the remote functional co-fluctuations (B. B. Biswal et al., 2010; Friston, 2011). Previous functional connectivity methods have typically assumed stationarity that the time series do not change their characteristics over time. However, the brain is indeed a complex system featured by the dynamic functional brain connectome (Zalesky et al., 2014). Recent evidence suggests that functional interactions between different brain regions and networks vary with time. The sliding-window based dynamic functional connectivity (dFC) is intensively used to measure the dynamic interaction between two regions (Hutchison et al., 2013; Preti et al., 2017), which has been shown to link with individual differences (Fong et al., 2019), task performances and disease alterations (Gonzalez-Castillo & Bandettini, 2018).

Despite the limitation of the sliding-window approach, the dynamic characteristics of the functional brain connectome at the macroscopic level support that the brain has multiple functional recurring states (Zalesky et al., 2014). There are several methods for dynamic brain state detection. Combining the sliding-window dFC with the clustering method, Allen and colleagues identified several FC states, as an intermediate scale between static and instant FC underlying small short-time tasks, which can reallocate and integrate attentional and executive resources (Allen et al., 2014). Besides, based on the assumption of temporal independence, Smith and colleagues used temporal ICA to identify temporal function modes (TFM), which represent unique brain activation patterns (Smith et al., 2012). Different from the dFC-clustering and TFM which assign each time point to one single state, the approach of the hidden Markov model (HMM) could identify a mixture of the brain states with a given probability at each time point, by assuming the transitions between states should follow a Markov process (Vidaurre et al., 2016). Vidaurre et al. used HMM in resting-state fMRI, and they found two hierarchical metastates that represent higher-order cognition and sensorimotor systems (Vidaurre et al., 2017).

The coactivation pattern (CAP) analysis is a data-driven method to detect the functional brain states in a single volume level (Liu et al., 2013; Liu & Duyn, 2013), which originates from the point process analysis (Tagliazucchi et al., 2012). Rather than capturing the dFC configurations as brain states, it is simple and straightforward that, for each frame of the data, the spatial coactivation patterns represent a specific whole-brain activation configuration to deal with the real-time task at that time, and that different frames which share the same spatial patterns are regarded as the same CAP state (Figure 1A). As a data-driven method, CAP analysis relies on very few mathematical presumptions, and is free of the confounding influence from the sliding window length. Therefore, CAP analysis has increasingly been used to study the abnormal network dynamics in depression (Kaiser et al., 2019), Alzheimer’s disease (Kaiser et al., 2019; Ma et al., 2020) and task fMRI (Freitas et al., 2020).

**Figure 1.**
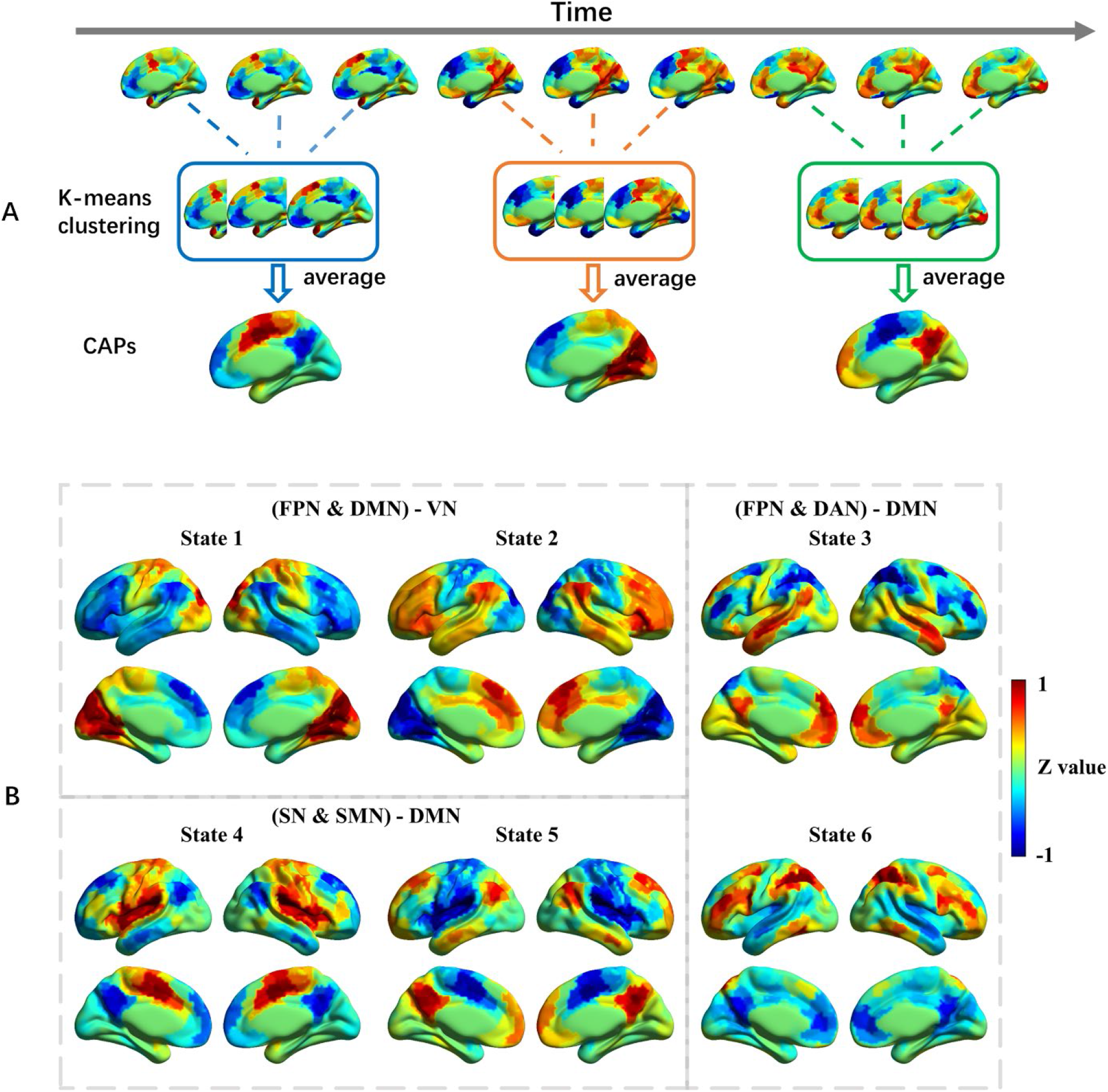
A) An illustration for the CAP analysis. The normalized spatial map for each volume was input for the k-means clustering, to identify volumes with similar coactivation patterns, and then average them to generate the CAPs. B) Six CAP states were identified by CAP analysis based on the primary configuration (WuXi cohort, resting-state fMRI preprocessing with GSR and 408 ROIs for the network construction). These brain states were normalized at the group level, and the value is the z-statistic value. Red color indicates a relatively stronger activation, while blue color indicates a relatively stronger deactivation. State 1 was mainly related to deactivated FPN, DMN and activated VN, and the opposite is true for State 2; State 3 was mainly characterized by activated DMN and deactivated FPN and DAN, and the opposite is true for State 6; State 4 was mainly characterized by deactivated DMN and activated SN and SMN, and the opposite is true for State 5. Abbreviations: DAN, dorsal attention network; DMN, default mode network; FPN, fronto-parietal network; SN, salience network; SMN, somatomotor network; VN, visual network.

The reproducibility is crucial for analytical methods in fMRI studies (Botvinik-Nezer et al., 2020; Eklund et al., 2016; Zuo et al., 2019). A lot of analytic flexibility exists in fMRI studies (Vergara et al., 2017), such as different preprocessing pipelines (Shirer et al., 2015), different software and toolboxes (Bowring et al., 2019). It is necessary to clarify the effects of varied settings in neuroimaging analyses (Aurich et al., 2015; Strother, 2006; Vergara et al., 2017) and to establish a standard and robust methodological pipeline (Esteban et al., 2019). However, it still remains unknown about the analytic flexibility and reproducibility for CAP analysis.

Schizophrenia is a psychiatric disorder with complex structural and functional brain alterations, characterized by abnormal connectome and functional dynamics (Collin et al., 2016; Fornito et al., 2012; Hunt et al., 2017). Particularly, the triple-network (V. Menon, 2011), including the default-mode network (DMN), fronto-parietal network central (FPN) or executive network (CEN), and salience network (SN), is postulated as a critical core of network dysfunction for understanding the neuropathological mechanism of psychiatric disorders including schizophrenia (Manoliu et al., 2014; V. Menon, 2011). Recently, Supekar and colleagues found that reduced dynamic interactions among the SN, CEN and DMN may substrate neurobiological signatures of schizophrenia (Supekar et al., 2019). However, little is known about the aberrant dynamic characteristics in schizophrenia concerning triple networks as well as other parts of the whole brain.

This study aimed to first investigate the reproducibility and generalizability of the CAP, and then utilize the robust analytical approach to study the aberrant dynamic state transition in schizophrenia. To achieve this goal, key methodological aspects were carefully evaluated for the robustness of CAP, including different preprocessing pipelines, ROI numbers for network construction, cluster numbers, and cohorts. Then, reliable dynamic states in the functional brain coactivation patterns were identified and further employed to compare the temporal dynamic characteristics between schizophrenia patients (SZ) and healthy controls (HC) in three independent data cohorts. Next, the associations between aberrant dynamic state transition and clinical symptom severity were explored.

## 2. Materials and Methods

### 2.1 Participants

To explore the reproducibility and generalizability of this study, three cohorts (WuXi, COBRE and UCLA) were analyzed. The WuXi cohort was used as the primary cohort, the other two open-access cohorts were treated as verification and their detailed participant information was described in the Supplementary material.

For the primary cohort (WuXi), all subjects were scanned by a structural MRI and resting-state functional MRI on a 3.0-Tesla Magnetom TIM Trio (Siemens Medical System) at the Department of Medical Imaging, Wuxi People’s Hospital, Nanjing Medical University. Foam pads were used to reduce head motion and scanner noise. Before the scan, the subjects were instructed to keep their eyes closed, relax but not fall asleep, and move as little as possible during data acquisition. After excluding subjects with large head motion, 69 SZ subjects and 97 HC subjects remained for the current study. Positive and Negative Syndrome Scale (PANSS) was used to measure the psychiatric symptoms of SZ patients.

Framewise displacement (FD) was calculated from the resting-state fMRI data to measure head motion (Di & Biswal, 2015). Subjects were excluded if their maximum translation or rotation FD were greater than 2 mm or 2°. The k-means clustering was performed in all 97 HC subjects. For the group comparisons between SZ and HC, only age- and gender-matched HC subjects were analyzed. The age- and gender-matched demographic information for the three cohorts were provided in Table 1. The information for all HC subjects used in the coactivation patterns generation can be found in the Supplementary material.

**Table 1.** The demographic information for the three cohorts

|  | HC | SZ | P value |
| --- | --- | --- | --- |
| **WuXi cohort ($n = 138$)** |  |  |  |
| Num of subjects | 69 | 69 |  |
| Age [years] | $45.84 \pm 11.89$ | $46.06 \pm 10.96$ | 0.9112<sup>a)</sup> |
| Gender (Male / Female) | 35 / 34 | 35 / 34 | 1<sup>b)</sup> |
| Disease duration | - | $19.84 \pm 10.96$ | - |
| PANSS positive | - | $20.06 \pm 4.59$ | - |
| PANSS negative | - | $23.78 \pm 3.84$ | - |
| PNASS general | - | $41.67 \pm 5.27$ | - |
| PNASS total | - | $85.51 \pm 9.50$ | - |
| **COBRE cohort ($n = 108$)** |  |  |  |
| Num of subjects | 54 | 54 |  |
| Age [years] | $37.22 \pm 12.48$ | $37.80 \pm 14.13$ | 0.8234<sup>a)</sup> |
| Gender (Male / Female) | 43 / 11 | 43 / 11 | 1<sup>b)</sup> |
| Disease duration | - | $15.19 \pm 12.46$ | - |
| PANSS positive | - | $14.13 \pm 4.29$ | - |
| PANSS negative | - | $14.46 \pm 5.10$ | - |
| PNASS general | - | $28.57 \pm 8.40$ | - |
| PNASS total | - | $57.17 \pm 13.52$ | - |
| **UCLA cohort ($n = 90$)** |  |  |  |
| Num of subjects | 45 | 45 |  |
| Age [years] | $36.73 \pm 8.65$ | $37.00 \pm 8.75$ | 0.8973<sup>a)</sup> |
| Gender (Male / Female) | 33 / 12 | 33 / 12 | 1<sup>b)</sup> |
| BPRS | - | $50.40 \pm 14.08$ | - |
| SAPS | - | $28.89 \pm 18.49$ | - |
| SANS | - | $35.09 \pm 18.55$ | - |

Data are expressed as mean ± SD (SD: standard deviation).
Abbreviations: BPRS, Brief Psychiatric Rating Scale; SANS, Scale for the Assessment of Negative Symptoms; SAPS, Scale for the Assessment of Positive Symptoms; PANSS, Positive and Negative Syndrome Scale.
<sup>a)</sup> two-sample t-test; <sup>b)</sup> chi-square cross-table test.

### 2.2 fMRI Data Acquisition

for the primary cohort (WuXi), the resting-state scans were acquired using a single-shot gradient-echo echo-planar-imaging sequence (Tian et al., 2016) with the following parameters: TR = 2000 ms, TE = 30 ms, slice number = 33, slice thickness = 4 mm, flip angle = 90°, matrix size = 64 × 64, FOV = 220 mm, voxel size = 3.4 × 3.4 × 4 mm^3^, and volume number = 240. Three-dimensional T1-weighted images were acquired by employing a 3D-MPRAGE sequence with the following parameters: TR = 2530 ms, TE = 3.44 ms, flip angle = 7°, matrix size = 256 × 256, slice number = 192, slice thickness = 1 mm, FOV = 256 mm, and voxel size = 1 × 1 × 1 mm^3^. The fMRI data acquisition parameters for the other two cohorts were described in the Supplementary material.

### 2.3 fMRI Data Preprocessing

The resting-state fMRI data were preprocessed using DPABI (http://rfmri.org/dpabi), and the preprocessing steps were followed: 1) Remove the first 2 time points for the UCLA and COBRE dataset, and remove the first 5 time points for the WuXi cohort; 2) Realignment; 3) Coregisteration of T1 image to functional image; 4) T1 segmentation by DARTEL; 5) Normalization of the functional images by T1 DARTEL; 6) Nuisance regression, including 24 head motion parameters, mean white matter (WM) and mean cerebrospinal fluid (CSF) signal, both with and without global signal regression (GSR); 7) Detrend; 8) Band-pass filtering, from 0.01 ~ 0.08 Hz; 9) Smoothing with an 8 mm FWHM kernel.

To evaluate the effect of preprocessing steps, the resting-state fMRI data was also preprocessed using a standard task fMRI data preprocessing pipeline, which is similar to the above steps but without nuisance regression and filtering.

The BOLD signal for the preprocessed resting-state fMRI data was extracted from 408 ROIs separately. The 408 ROIs were consist of 400 cortical regions from Yeo’s 7 network parcellation (Schaefer et al., 2018) and 8 subcortical regions (bilateral caudate nucleus, putamen, globus pallidus and amygdala) from the AAL template (Tzourio-Mazoyer et al., 2002). The 400 cortical regions are allocated to 7 networks including the visual network (VN), somatomotor network (SMN), dorsal attention network (DAN), ventral attention network (VAN), limbic network, fronto-parietal network (FPN) and default mode network (DMN) (Yeo et al., 2011). The 7 networks have parcellations with different spatial scales from 100 to 1000, and 400 was mainly used in this study because the average voxel size for the 400 ROIs is comparable to the average voxel size of the 8 subcortical regions from the AAL template.

### 2.4 Coactivation Pattern Analysis

Coactivation pattern (CAP) analysis is a data-driven method that identifies recurring states across time points with similar whole-brain coactivation patterns. In this study, the CAP analysis was performed using home-made scripts in MATLAB (https://www.mathworks.com/).

First, to represent the relative activation magnitude changes in the 408 ROIs, each time series were normalized using a z-score. For each subject *i*, a two-dimensional normalized BOLD matrix ***X_i_*** (*T*× 408) was obtained, where T is the number of time points and 408 is the ROI number.

Next, all HC subjects’ normalized BOLD metrics ***X_HC_i_*** (*T* × 408) were concatenated to obtain the ***X_HC_*** (*T_HC_* × 408, *T_HC_* = *T*× *N*), where *N* is the sample size of all HC subjects. Then, k-means clustering was performed to identify similar coactivation patterns across all volumes from all HC subjects, and the distance between two volumes was calculated by subtracting their Pearson correlation coefficient from one. The cluster number K was selected from 2 to 21 with a step length of 1. The clustering algorithm was repeated 100 times with a new initial cluster centroid for each K value, and the results with the lowest within-cluster sums of point-to-centroid distances were used. Frames assigned to the same CAP state were averaged and divided by the within-cluster standard deviation to generate the normalized CAP maps (Z-maps) at the group level (Figure 1A).

The clustering patterns obtained from all HC subjects were then applied to each SZ subject. Specifically, each frame from the ***X_SZ_i_*** (*T* × 408) was extracted, which is a 1 × 408 vector representing the whole-brain coactivation level at that time point. Then, the spatial similarity between each frame and each normalized CAP map was calculated using the Pearson correlation, and the frame was assigned to the CAP with the largest spatial similarity.

The silhouette score (Rousseeuw, 1987) was calculated to evaluate the clustering results for different K values. As shown in Figure S1, Supplementary material, the silhouette score was monotonically decreasing with the increase of K. Then, the elbow criterion was considered to determine the number of clusters. While one issue is that the time points of the three cohorts are limited, if the cluster number is too large, then each CAP state would only account for a few seconds through the entire scan. Therefore, 6 clusters were mainly analyzed and reported in the manuscript as a trade-off, and one recent paper also used 6 clusters for the CAP analysis (Zhang et al., 2020).

### 2.5 State Temporal Dynamics Measures / CAP Metrics

To evaluate the dynamic properties within and between CAP states, four dynamic measures (CAP metrics) were calculated at the individual level: 1) **Fraction of time** is defined as the proportion of total volumes spent in one CAP state over the whole time series; 2) **Persistence** is the average time spent in one state before transferring to another state, and it describes the mean volume-to-volume maintenance of one CAP state; 3) **Number of states (Counts)** is how many times one state occurred during the whole scan; and 4) **Transition probability matrix** is the probability that one volume within State A transfers to the next volume belonging to State B, with a non-zero diagonal as the volume within State A could still stay within State A for the next volume.

### 2.6 Transition Probability and Spatial Similarity between States

The relationship between the spatial similarity and transition probability between two states was measured in all HC subjects. The spatial similarity between different brain states was calculated using the Pearson correlation. Before measuring the relationship between the two metrics, the symmetry of the transition probability was first examined. In detail, the transition probability from State A to State B was paired with the transition probability from State B to State A, and the Pearson correlation was calculated between all pairs to test the symmetry level. The transition probability metrics were then averaged on the group level and symmetrized. Finally, the relationship between the spatial similarity and transition probability was measured using the Pearson correlation. The diagonal values were not analyzed in this part.

### 2.7 Reproducibility Analysis

Recently, neuroimaging studies have drawn more attention to reproducibility. In this study, the CAPs’ spatial and temporal reproducibility were considered from four aspects. The first aspect is to consider the effects of different data preprocessing steps, including two standard resting-state fMRI preprocessing pipelines (with and without GSR) and one classic task fMRI preprocessing pipeline. Then, the effects of different spatial resolutions of the template were assessed. In detail, 100, 200, 400 and 1000 ROIs from Yeo’s 7 network parcellations (Schaefer et al., 2018) plus the 8 subcortical regions from the AAL atlas (Tzourio-Mazoyer et al., 2002) were tested. Different K values were also compared to see whether the CAP states gradually change with the increase of cluster number. Finally, the CAP analysis was performed in three independent cohorts separately to detect the site effect.

Furthermore, to verify the generalizability of this study, whether the results obtained by one cohort can be directly replicated in other cohorts, we applied the CAPs generated from the WuXi cohort to the other two cohorts, then the spatial maps and temporal dynamics among CAP states were compared.

The spatial reproducibility was assessed by calculating the Pearson correlations between CAPs’ spatial maps under different conditions, and the temporal dynamics were then compared between spatial matched CAPs.

### 2.8 Statistical Analysis

For group comparisons, only age- and gender-matched HC and SZ subjects were analyzed. Age was compared between SZ and HC by a two-sample t-test, and a chi-square cross-table test was used to test the gender difference. As for the CAP metrics, two-sample t-tests were performed with age and gender as covariates. FDR correction (q = 0.05) was used to correct for the multiple comparisons.

The relationship between CAP metrics and behavioral measures, such as disease symptoms and disease duration, were tested using partial correlation with age and gender as covariates. The state temporal dynamics and behavioral measures were first normalized using z-score, and subjects with large deviation (Z > 3) were excluded in the correlation analysis as the outlier. FDR correction (q = 0.05) was used to correct for the multiple comparisons.

## 3. Results

### 3.1. Demographics and Questionnaires

No group differences of age or gender were found between the SZ and HC in all three cohorts, and the detailed group characteristics are given in Table 1.

### 3.2. Reliable CAP identification and Dynamic Characteristics

This study tested different methodological combinations on three independent data cohorts, such as preprocessing pipelines, ROI numbers and cluster numbers. Below we reported the primary results of six reliable CAP states, which are based on the WuXi cohort, using the resting-state preprocessing with global signal regression (GSR) and 408 ROIs. The other validation results are provided in the Supplementary material.

#### 3.2.1. Coactivation Patterns and Brain States

The coactivation patterns were generated from all HC subjects using temporal k-means clustering, and six CAP states were finally identified (Figure 1B), after the search for optimal cluster number (see details in the Supplementary material). Among the six CAP states, brain regions that belong to the same functional network, such as the default mode network (DMN), fronto-parietal network (FPN), and salience network (SN), tend to be activated or deactivated simultaneously. One interesting phenomenon is that the six CAP states were grouped into three pairs with opposite spatial coactivation patterns. For example, State 1 and 2 grouped together – State 2 was mainly related with the activated FPN, DMN (without posterior cingulate cortex and precuneus) and deactivated visual network (VN), while State 1 had the opposite spatial pattern. Since each CAP state had certain brain networks with relatively stronger activation or deactivation than the other networks of the whole brain, we found that not only triple networks but also primary (VN, SMN) and higher-order networks (DAN) were identified in the dominant CAP states.

#### 3.2.2. Transition Probability and Spatial Similarity between States

As shown in the diagonal of the transition probability matrix (Figure 2A), the temporal activity was dominated by the identified CAP states (more than 60% of the time), and the between-state alteration remained low transition probability (1% to 11 % of the time). Comparing Figure 2A and Figure 2B, it is clear that the transition probability between brain states with strong anti-correlated spatial coactivation was close to zero. Taking State 4 and State 5 for an example (Figure 2C), their coactivation patterns were opposite (spatial similarity r = - 0.99), their transition probability from State 4 to State 5 was 0.0081, and from State 5 to State 4 was 0.0103. Despite the small discrepancy in bi-directional transition probability, the symmetry in the transition probability matrix is pronounced. Figure 2D showed a significant positive correlation (r = 0.8479, p < 0.0001) between the transition probability from State A to State B and transition probability from State B to State A. Finally, the relationship between the transition probability and spatial similarity metrics was evaluated. As shown in Figure 2E, the transition probability between two CAP states was highly correlated with their spatial similarity (r = 0.9817, p < 0.0001).

**Figure 2.**
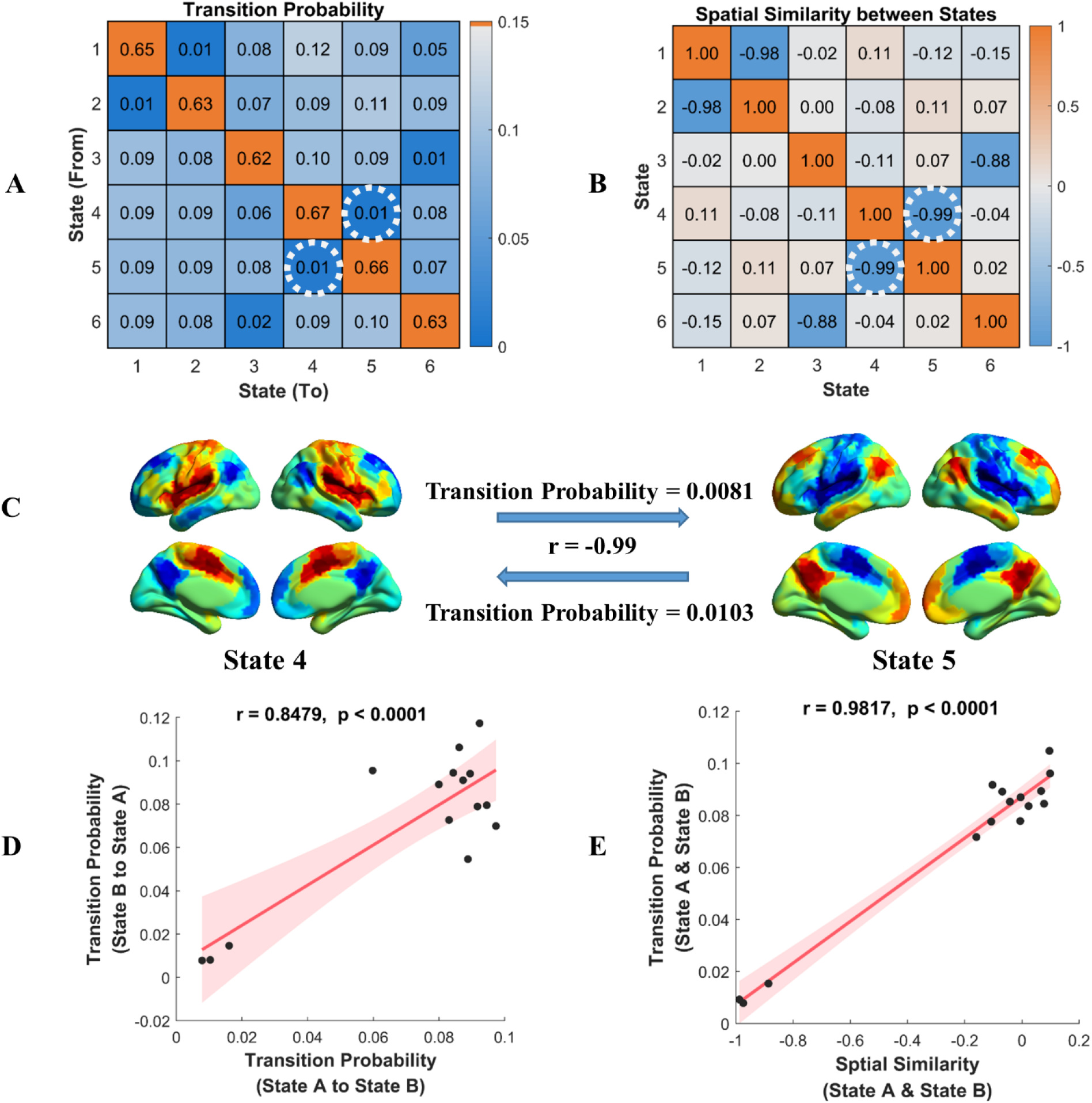
The relationship between CAP state transition probability and their spatial similarity. A) The group average transition probability matrix in all HC subjects. B) The spatial similarity between six CAP states, measured by the Pearson correlation. C) The transition probability between State 4 and State 5, and their spatial similarity, the values were shown in the white dashed circles in Figure 2A and Figure 2B. D) The correlation between transition probability from State A to State B and transition probability from State B to State A. The shadow represents the 95% confidence interval. E) The correlation between the symmetrized transition probability between State A and State B and their spatial similarity. The shadow represents the 95% confidence interval.

#### 3.2.3. Spatial Reproducibility Evaluation

As shown in Figure 3, each row or column showed two pairs with high spatial similarity, which suggest reproducible CAP states and their groups. It indicated that good spatial reproducibility within the HC group remained across different preprocessing pipelines and ROI numbers for network construction.

**Figure 3.**
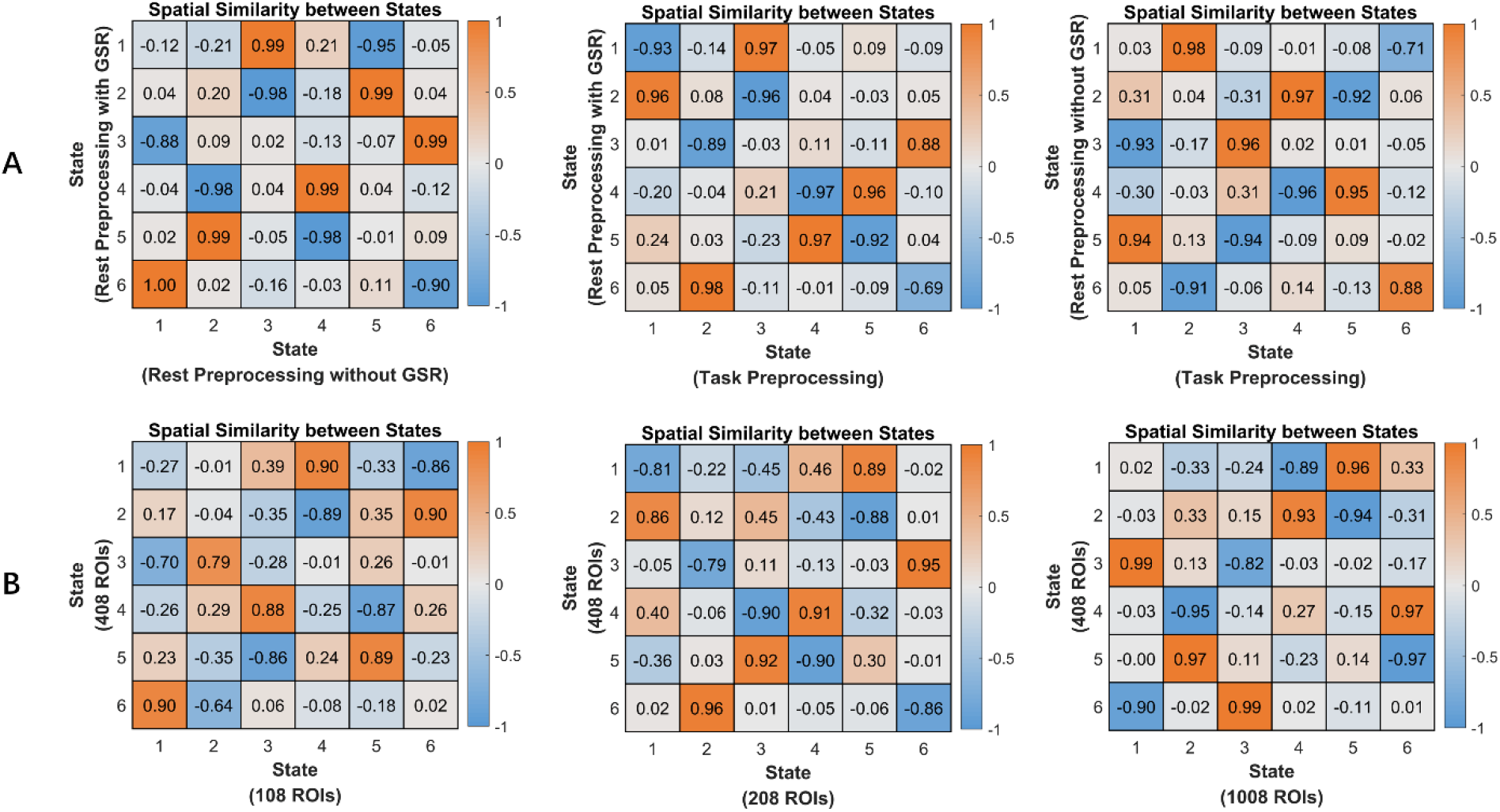
The CAP spatial similarity between states under A) different preprocessing pipelines and B) different ROI numbers. The spatial similarity was measured by the Pearson correction.

Moreover, the validation analysis for different cluster number K also demonstrated good spatial reproducibility, although more CAP states were separated with the increase of K. For example, as shown in Figure 6, when K increased from 6 to 8, four states (State 3 to State 6) remained unchanged, their one-to-one correspondence spatial similarity was larger than 0.9, and the other two states (State 1 and State 2) were subdivided into four states.

To evaluate the reproducibility and generalizability of identified CAP states, the clustering results from the WuXi cohort were applied to all subjects from the COBRE cohort and UCLA cohort. The diagonal of metrics in Figure 7A showed the high spatial similarity of each CAP state between the WuXi cohort and the other two cohorts.

#### 3.2.4. Temporal Reproducibility Evaluation

To verify the temporal reproducibility within the HC group more straightforwardly, all states were relabeled to group corresponding states together. As shown in Figure 4, the absolute values for the CAP metrics across different preprocessing pipelines, ROI numbers and cohorts were evaluated in HC. As for the preprocessing pipeline, rest preprocessing with and without GSR showed consistent results across all the three CAP metrics, while task preprocessing showed shorter persistence and more counts. All three CAP metrics were not sensitive to the ROI number. Different cohorts showed a consistent fraction of time and persistence, and the WuXi cohort exhibited more counts than the other two cohorts.

**Figure 4.**
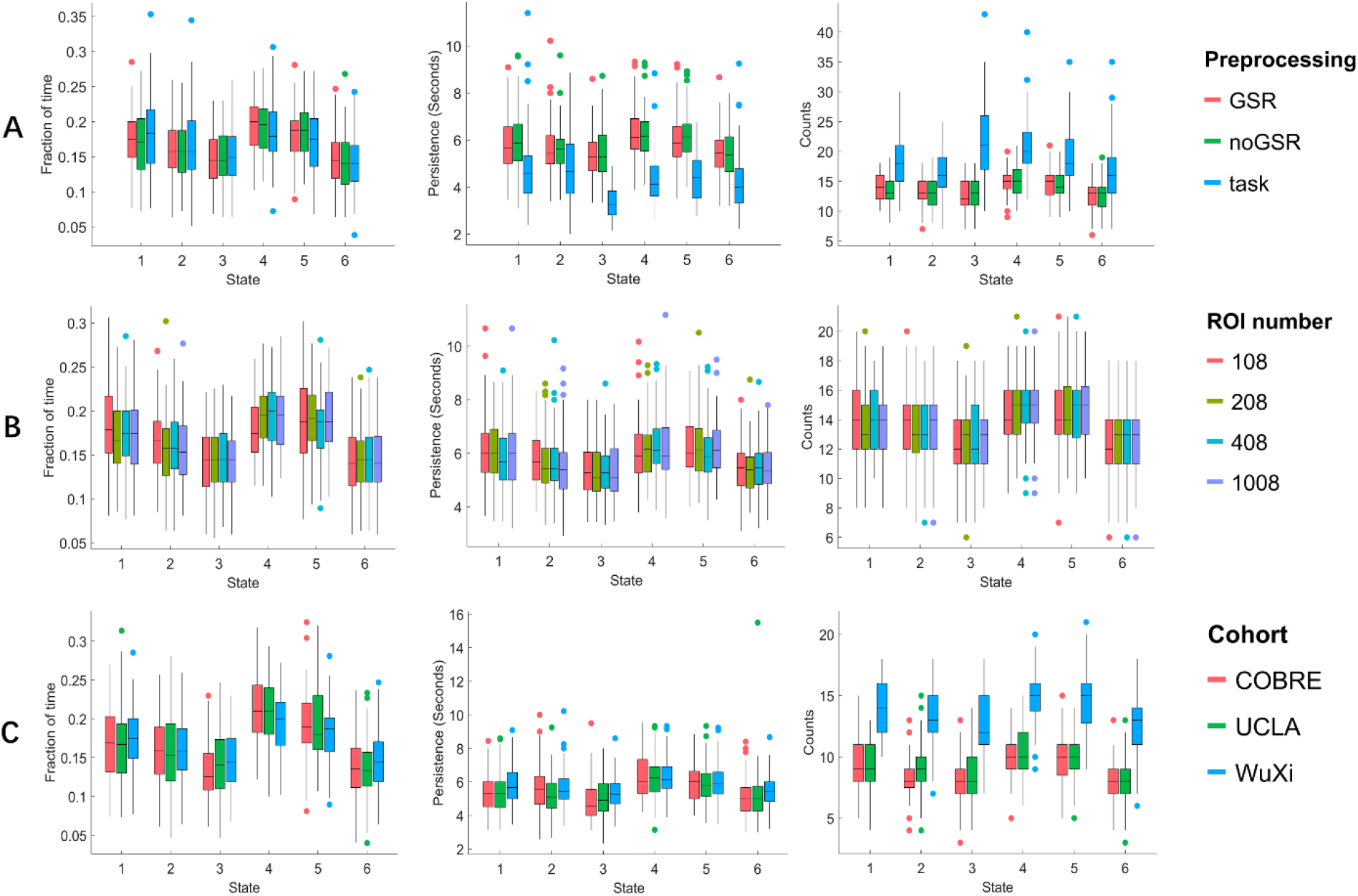
The CAP metrics reproducibility within the HC group under different A) preprocessing pipelines, B) ROI numbers and C) cohorts.

### 3.3. Aberrant and Reproducible State Dynamics in Schizophrenia

#### 3.3.1. State Temporal Dynamic Differences between SZ and HC

The robust CAP analysis was applied to investigate the schizophrenia-related abnormalities in the CAP dynamic state transition across three independent data cohorts. The state temporal dynamics (CAP metrics) were compared between SZ patients and well-matched HC controls by using a two-sample t-test, with age and gender as covariates. The results of group comparisons were presented in Figure 5. As mentioned above, the six CAP states could be grouped into three pairs (State 1 and 2, State 3 and 6, State 4 and 5). The mean fraction of time of each state for SZ and HC groups was around 15% to 20%, and each state persisted for 5 to 6 seconds. For example, the pair of State 4 and State 5 occupied the highest fraction of time in both the SZ and HC groups, which shared opposite spatial coactivation patterns dominated by SN, SMN and DMN. Almost every state except State 6 showed significant temporal dynamic differences (P<0.05, FDR corrected). The group differences were similar within each pair. For instance, SZ patients showed less fraction of time in states characterized by FPN and DMN (State 1 and State 2), and more fraction of time in states characterized by SN and DMN (State 4 and State 5).

**Figure 5.**
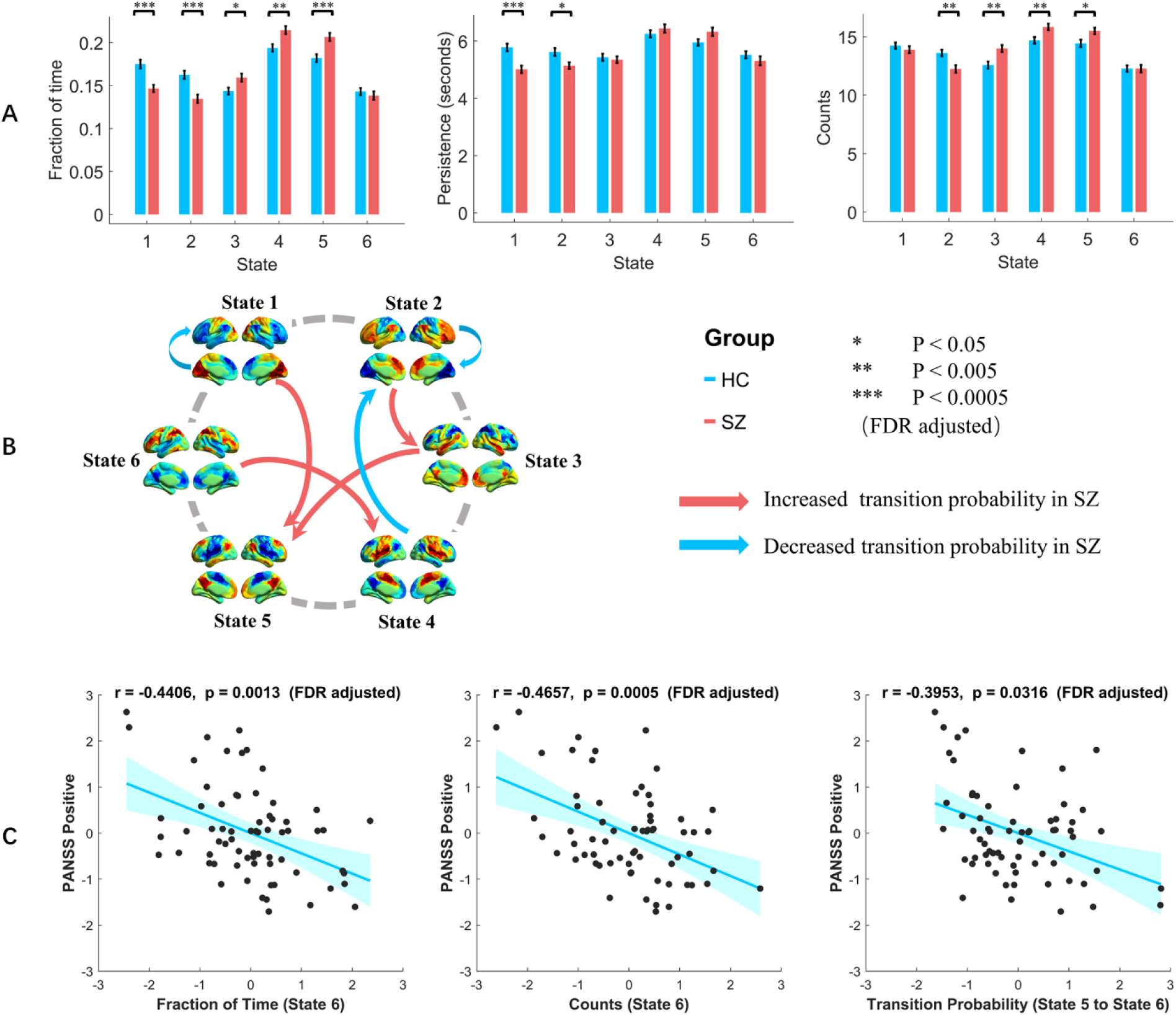
State temporal dynamic differences between SZ and HC. The group differences in A) fraction of time, persistence, counts and B) transition probability. Red bins are the SZ group and blue bins are the HC group. Two-sample t-tests were performed with age and gender as covariates. Error-bar is the standard error. * indicates p < 0.05, and ** indicates p < 0.005, and *** indicates p < 0.0005 with FDR correction. For the transition probability, the red arrow means higher transition probability in the SZ group, and vice versa for the blue arrow. C) Behavioral relevance with state temporal dynamics in SZ. The fraction of time of State 6 was negatively correlated with the positive PANSS score, r = −0.4406, p = 0.0013; The counts of State 6 was negatively correlated with the positive PANSS score, r = −0.4657, p = 0.0005; The transition probability from State 5 to State 6 was negatively correlated with the positive PANSS score, r = −0.3953, p = 0.0316 (all the p values were FDR adjusted). The shadow represents the 95% confidence interval. Abbreviations: PANSS, Positive and Negative Syndrome Scale.

Specifically, SZ patients showed a significantly reduced fraction of time and persistence in State 1 and 2, as well as reduced counts in State 2, compared with the HC group (Figure 5A). In State 4 and 5, SZ patients had significantly increased fraction of time and counts. Moreover, SZ patients showed a significantly increased fraction of time and counts in State 3, but not in State 6. As for the transition probability between CAP states, SZ patients showed lower transition probability from State 4 to State 2, and lower transition probability within State 1 and State 2 (Figure 5B). On the other hand, SZ patients showed higher transition probability from State 1 to State 5, State 2 to State 3, State 3 to State 5 and State 6 to State 4. The detailed statistic values for these CAP metrics were described in Table S2, Supplementary material.

#### 3.3.2. The relationships between State Temporal Dynamics and Clinical Data

The clinical relevance with state temporal dynamics was evaluated in the SZ group, using partial correlation with age and gender controlled. As shown in Figure 5C, after FDR correction, the following negative correlations between CAP metrics and positive PANSS score were found: the fraction of time of State 6 (r = −0.4406, p = 0.0013), the counts of State 6 (r = −0.4657, p = 0.0005), and the transition probability from State 5 to State 6 (r = −0.3953, p = 0.0316). In addition, the persistence of State 3 (r = −0.3388, p = 0.0323) was negatively correlated with the disease duration, the fraction of time of State 4 (r = 0.3556, p = 0.0203), and the persistence of State 5 (r = 0.3653, p = 0.0154) was positively correlated with the PANSS total score.

#### 3.3.3. Reproducible Group Differences between SZ and HC

The reproducibility of SZ patients’ dynamic alterations was also validated in this study, which confirmed good temporal reproducibility for the group differences. For instance, SZ showed more fraction of time in State 4 and State 5 and less fraction of time in State 1 and State 2. These results were consistent across different methodological pipelines (Figure S5 and Figure S8, Supplementary material). And for the unchanged states under K = 6 (State 5) and K = 8 (State 1), their temporal dynamic differences that SZ showed more fraction of time than HC were consistent as well (Figure 6B).

**Figure 6.**
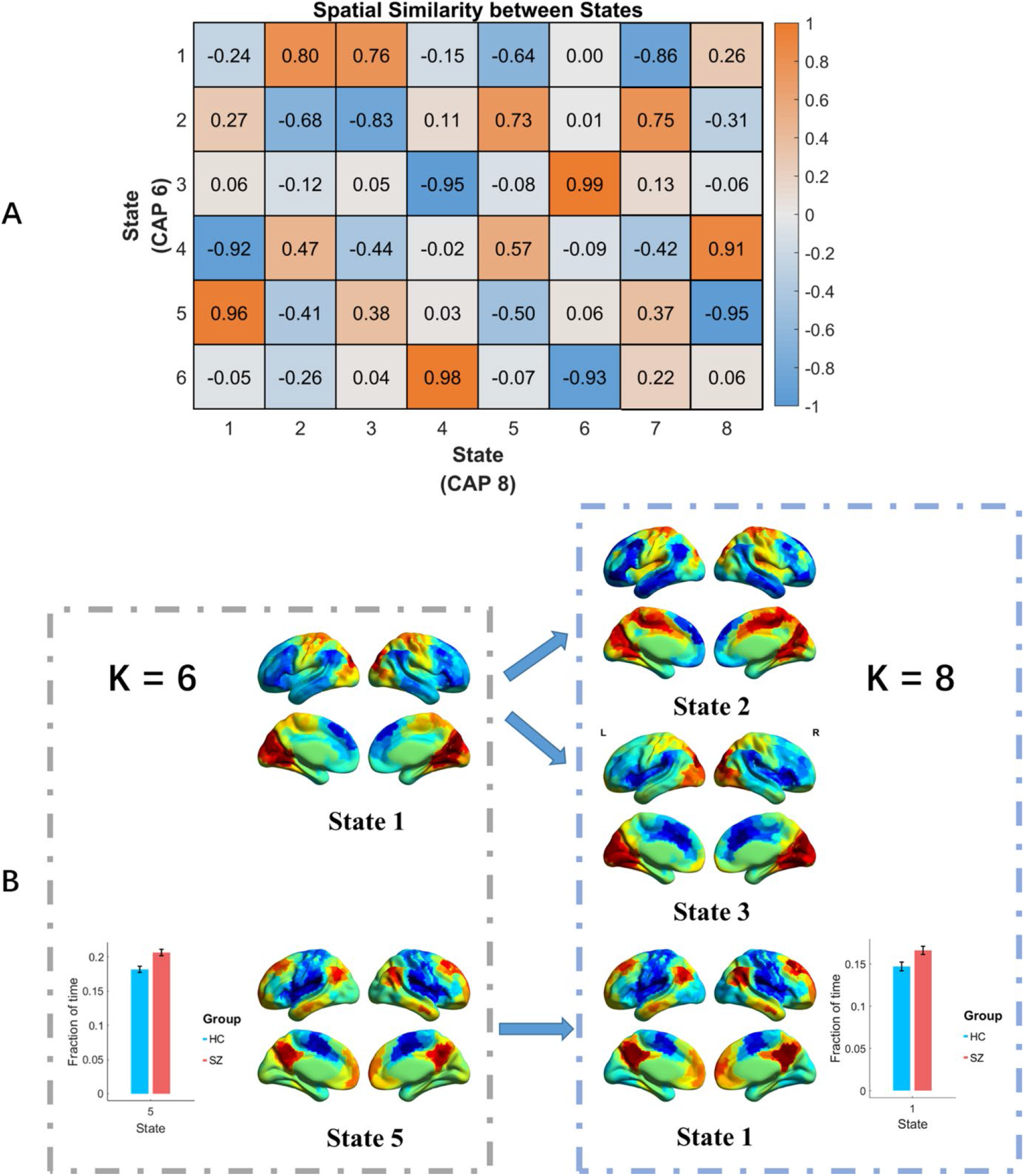
CAP analysis reproducibility with different cluster numbers. In this case, the configuration was WuXi cohort, rest preprocessing with GSR and 408 ROIs. A) The CAP spatial similarity between K = 6 and K = 8. B) The State 1 in K = 6 was divided into two states in K = 8, State 2 and State 3. State 5 in K = 6 remained in K = 8, which corresponds to State 1, and the SZ group also showed a consistent more fraction of time than the HC group. The error bar is the standard error.

Figure 7B showed the fraction of time differences between SZ and HC across the three cohorts. Their overall trend among the six CAP states was similar, particularly State 2, State 3 and State 5 showed consistent significant group differences in the WuXi cohort and COBRE cohort. Although the temporal dynamic differences obtained from the UCLA cohort were less similar compared with the other two cohorts, the absolute values still showed a consistent trend as presented in Figure 5C.

**Figure 7.**
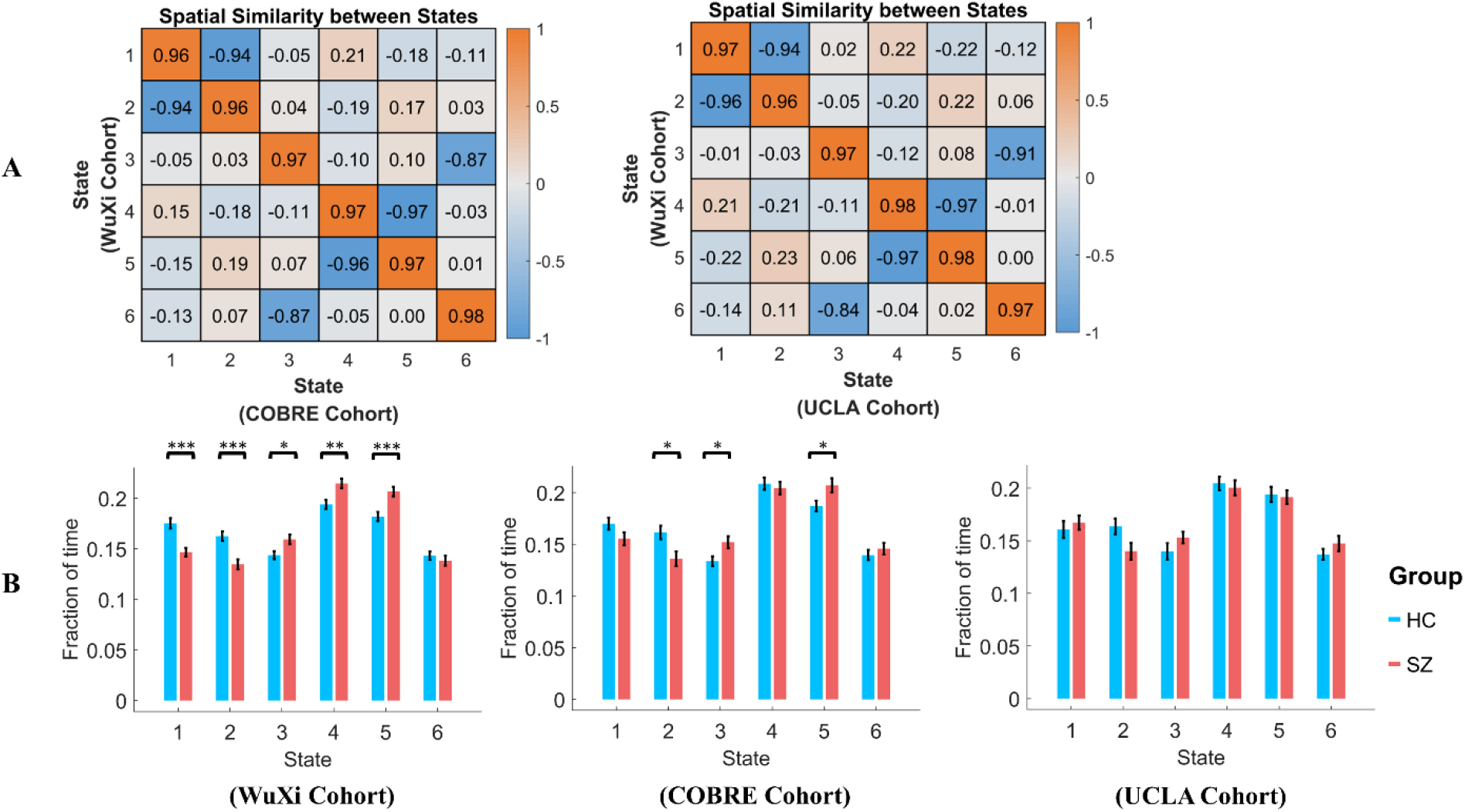
The generalizability of network dynamic measure differences between SZ and HC across the three independent cohorts. A) Applying the clustering results based on the WuXi cohort to the other two cohorts, their spatial similarity was measured by the Pearson correlation. B) The fraction of time differences between SZ and HC across the three cohorts. * indicates p < 0.05, ** indicates p < 0.005 and *** indicates p < 0.005 (FDR adjusted). The error bar is the standard error.

In addition, the repeatability for different cohorts was validated. Rather than using the CAP maps from one cohort to the other cohorts, the CAP analysis was independently performed for the COBRE and UCLA cohort, and the spatial and temporal results were compared. Although the repeatability for different cohorts was relatively weaker, considerable spatial overlaps were identified across the three cohorts. More details were described in Figure S11 to Figure S13, Supplementary material.

## 4. Discussion

In this study, we first identified the characteristic and reliable states and transitions of functional brain networks in the healthy adults, and then investigated the schizophrenia-related aberrant state dynamics, based on the coactivation pattern analysis and three independent cohorts. Healthy and patient cohorts achieved robust results across different methodological pipelines. Our results revealed six reliable coactivation states of functional brain networks, which were constituted by typical resting-state networks including triple networks as well as primary and other higher-order networks. The principle of spontaneous state transitions inferred the higher spatial similarity, the higher transferring probability between the separable coactivation states. Patients with schizophrenia showed reproducible evidence of aberrant coactivation patterns dynamics, particularly the state dominance such as the fraction of time was altered and associated with positive symptoms in the patients. Together, our study confirms the reproducibility and generalizability of CAP analysis, which could provide meaningful information about the network dysfunction and neuropathological mechanisms in psychiatric disorders.

### 4.1. Coactivation Patterns for Brain States

The CAP analysis is based on the temporal k-means clustering of whole-brain functional activities, which identifies a group of spatial maps with similar whole-brain coactivation patterns across the whole scan. Motivated by the idea of PPA (Tagliazucchi et al., 2012), Liu et al. found that the DMN can be simply identified by averaging multiple distinct coactivations or co-deactivation patterns at different time points (Liu & Duyn, 2013). Yeo’s 7 network parcellation was used in this study, and brain regions belonging to the same network tended to be activated or deactivated together (Figure 2). This result supports the intrinsic relationship between brain regions within the same functional network (B. B. Biswal et al., 2010; Calhoun & Adali, 2012), and these intrinsic networks can be simply extracted by averaging a few time points rather than using a more complex mathematical method such as ICA. Thus, while CAP states are derived from resting-state fMRI signals by using not pairwise functional connectivity but temporal clustering, which we feel at least partly represents the temporal dynamic characteristics of the whole-brain functional connectome (Zuo et al., 2012).

Previous ICA analyses used to divide the DMN network into anterior part and posterior part (B. B. Biswal et al., 2010; Zuo et al., 2010). Our results also found the posterior DMN (mainly includes the precuneus/posterior cingulate cortex and angular gyrus) and the anterior DMN (mainly includes the medial prefrontal cortex) were activated at different level across states. The posterior DMN tends to be related to the SN and SMN (State 4 and 5), while the anterior DMN was associated with FPN and DAN (State 3 and 6). Besides, three pairs of states were identified with opposite coactivation patterns. For instance, when DMN was activated, the FPN and DAN were deactivated (State 3), and vice versa is true for State 6. The phenomenon of opposite CAP pairs has also been found in previous studies (Huang et al., 2020; Janes et al., 2020; Zhang et al., 2020), suggesting these regions tend to be activated in an opposite manner that the activation of region A would suppress the activity of another region B, and vice versa.

The DMN is known as the task-negative network. For State 3 to State 6, when DMN was activated, other task-positive networks such as FPN and SN were either not activated or deactivated. These results verified the anti-correlation between the task-positive network and the task-negative network (Fox et al., 2005; Power et al., 2011). Based on CAPs, Li and colleagues concatenated a set of task activation maps from the Human Connectome Project, and validated the robust anti-correlated functional network (DMN) across multiple tasks (Li et al., 2020). However, we found that the DMN was not always activated conversely compared with FPN. Specifically, the medial-prefrontal subsystem and temporal subsystem of DMN were co-deactivated with FPN when the visual network was activated, and vice versa is true (State 1 and State 2). This is consistent with recent findings that the DMN and FPN are coactivated when evaluating internal information (Beaty et al., 2016; Zhu et al., 2017) and involved in task preparation (Koshino et al., 2011).

### 4.2. Transition Probability and Spatial Similarity

Unlike the Pearson correlation matrix which is mathematically symmetric, the transition probability between State A and State B is not equal. Nevertheless, a significant positive correlation was obtained (r = 0.8479, p < 0.0001) between the transition probability pairs, suggesting the transition probability between two states is an approximation (Figure 4A). In addition, the transition probability between two states was significantly correlated with their spatial similarity (r = 0.9817, p < 0.0001), which suggests that one state would transfer to the other state with a higher probability. This positive association was also found by a previous study based on the hidden Markov model (Vidaurre et al., 2017), as the brain should activate continuously, it is less likely that one CAP state would directly change to another state with opposite whole-brain coactivation configuration without any intermediate state.

### 4.3. Reproducibility in CAPs Analysis and Results

In this study, we considered the reproducibility of our analyses from several aspects, including preprocessing pipeline, ROI number, cluster number and cohort, and they showed consistent results.

For CAP analysis based on resting-state fMRI data, currently there is no standard preprocessing pipeline. Some studies used the common task preprocessing pipeline, which mainly includes realignment, spatial normalization and smoothing (Kaiser et al., 2019; Karahanoglu & Van De Ville, 2015). More studies used the standard resting-state preprocessing pipeline, which has additional steps such as nuisance signal regression (WM, CSF) and temporal filtering, with and without GSR (Karahanoglu & Van De Ville, 2015; Liu et al., 2013; Liu & Duyn, 2013; Ma et al., 2020). We used both the task and resting-state preprocessing pipelines (with and without GSR) in this study. As shown in Figure 3 and Figure 4, similar coactivation patterns and temporal dynamics were obtained for different preprocessing pipelines, which suggests the preprocessing has little effect on the whole-brain coactivation patterns. Besides, as Figure 4A showed, task preprocessing has shorter persistence and more counts. The reason could be that the high-frequency noise (Chen & Glover, 2015) was not filtered during the task preprocessing, which causes frequent fluctuations and shorter persistence.

Both voxel-level (Liu et al., 2013) and ROI-level (Janes et al., 2020) were studied in previous CAP analysis studies. Using ROI could reduce the dimension and save a lot of time and computational resources (Chen et al., 2015), while it could also decrease the spatial resolution and ignore spatial details. We chose different ROI numbers from 100, 200, 400 to 1000 to represent multiple levels of ROI size. As shown in Figure 3 and Figure 4, the coactivation patterns showed highly spatial and temporal dynamics consistency, suggesting that the CAP analysis is not sensitive to the spatial resolution.

As for the cluster number, when increasing the cluster number, the spatial and temporal properties change continuously. For instance, when K increased from 6 to 8, four states remained the overall spatial coactivation patterns, and their temporal dynamics were also unchanged (Figure 6).

Besides the analytic variations, we also validated the results obtained from the three independent cohorts. Although the spatial consistency of coactivation patterns between cohorts was less than that of different preprocessing steps, there were still considerable spatial overlaps. To further verify the generalizability of our findings, we mapped the CAP maps obtained by the WuXi cohort to the other two cohorts, and the group temporal dynamic differences between SZ and HC were similar across cohorts (Figure 7B). In conclusion, our study suggested there was considerable reproducibility across different analytic variations and cohorts.

### 4.4. State Temporal Dynamics Abnormalities in Schizophrenia and Their Reproducibility

Using the robust CAP spatial maps, the state temporal dynamics in terms of fraction of time, persistence, counts, and transition probability were calculated and compared between SZ and HC groups. Reproducible and aberrant state temporal dynamic was found in schizphrenia patients concerning different methodological pipelines or cohorts (Figure 5 - 7). Most CAP states demonstrated aberrant dynamic characteristics, suggesting schizophrenia-related network dysfunction is widespread over the whole brain, which is consistent with the accumulating evidence that schizophrenia is characterized by whole-brain network dysfunction (Adhikari et al., 2019; Collin et al., 2016; Fornito et al., 2012; Kambeitz et al., 2016; Venkataraman et al., 2012). Previous fMRI studies reported that schizophrenia patients showed distributed alterations in the dynamic functional connectivity (Du et al., 2018), dynamic brain activity (Fu et al., 2018) and dynamic state (Allen et al., 2014; Damaraju et al., 2014; Fu et al., 2020; Mennigen et al., 2018; Rashid et al., 2014). A recent resting-state fMRI study revealed dysregulated brain dynamics, i.e., reduced, less persistent, and more variable between-network interactions among SN, FPN and DMN in schizophrenia (Supekar et al., 2019), which proves aberrant triple network saliency model of psychosis (V. J. W. P. Menon, 2020). Our findings extend the current understanding about schizophrenia-related dynamic abnormalities in such manner that aberrant state temporal dynamics in schizophrenia is associated with not only triple networks but also part of primary (VN, SMN) and higher-order networks (DAN).

First, we found that SZ patients had insufficiently intensified activation and less inhibited deactivation in FPN-DMN state, but on the contrary for SN-DMN state, of which the transition probability changed significantly. It has been well documented that, the triple networks, involving FPN, SN and DMN, are the cores for higher cognition. Specifically, FPN is engaged in externally oriented attention during demanding cognitive tasks, SN is crucial in the process of salience mapping, and DMN is related to self-referential processes (V. Menon, 2011). Imaging findings based on triple network alterations have enhanced our understanding of the psychopathology in schizophrenia, depression and autism (Krishnadas et al., 2014; Manoliu et al., 2014; Nekovarova et al., 2014; Supekar et al., 2019; J. Wang et al., 2020). Recent meta-analysis confirms that the triple network might underlie the common network dysfunction across psychiatric disorders including schizophrenia (Sha et al., 2019). Although our findings highlight schizophrenia-related dynamic abnormalities centered in the triple network, in line with an earlier study (Supekar et al., 2019), our results further point out that the DMN was continuously activated across all states while SN and FPN were only involved with specific states, which may suggest the DMN play a crucial role in the state transitions and cross-network interactions within the triple networks.

Second, we found that the robust CAP states were also centered in the primary networks such as VN and SMN, and higher-order network such as DAN, which had substantially altered temporal state dynamics in schizophrenia patients. Interestingly, the VN, SMN, and DAN were recently identified across psychiatric disorders, by partial least squares which is a different data-driven approach from CAP analysis, as key parts underlying the general psychopathology, cognitive dysfunction, and impulsivity (Kebets et al., 2019). In this study, Kebets and colleagues found the latent components of whole-brain resting-state functional connectivity were robust, and particularly, SMN showed featured alterations in the static resting-state functional connectivity within and between networks. Our study provides consistent evidence for schizophrenia-related network dysfunction from a new perspective of CAP states and state transition. Furthermore, we reported that the dynamic characteristics of the FPN-DAN state (State 6) were negatively correlated with the PANSS positive scores, and those of SN-DMN state (State 4 and 5) were positively correlated with the PANSS total scores, consistent with previous evidence (Kindler et al., 2015; Manoliu et al., 2013; Pang et al., 2017; Rotarska-Jagiela et al., 2010; D. H. Wang et al., 2018). Importantly, the transition probability from the SN-DMN state (State 5) to the FPN-DAN state (State 6) was also negatively correlated with the positive PANSS scores. Notably, the group difference was not significant for State 6 (Figure 5A). This finding may suggest that the state transition is likely to alleviate the disease severity from the symptom positively-related state to the symptom negatively-related state, which might provide a potential intervention target for schizophrenia patients. Taken together, the reproducible abnormalities of state temporal dynamics identified in this study implicate that schizophrenia is associated with whole-brain functional network dysregulation and dynamic alterations.

### 4.5. Limitations

Although the k-means clustering has been widely used in fMRI data, currently there is no optimal criterion to determine the cluster number (Vergara et al., 2020). In this study, the volume numbers for the WuXi, COBRE and UCLA cohort are 240, 150 and 152 respectively, with the same TR. When we increased the cluster number in the CAP analysis, the average volumes allocated to each cluster (state) decreased, which might cause more variability for each cluster and reduce the clustering stability. In the COBRE and UCLA cohort, we tested the k-means clustering from 2 to 21, and for K = 21 there are only average 7 volumes for a single state, which had too limited temporal information. Therefore, we chose the K = 6 following the qualitative but arbitrary criteria in line with the prior study (Liu & Duyn, 2013).

## 5. Conclusion

In summary, functional brain states involved with specific coactivation patterns at different time points were obtained using coactivation pattern analysis. The spatial and temporal reproducibility of these CAPs was verified from multiple aspects, such as different preprocessing pipelines and independent cohorts. Moreover, the robust and aberrant temporal dynamics were identified in schizophrenia, associated with the severity of clinical symptoms. This study proved that the CAP analysis has good reproducibility and generalizability, which is useful to provide novel and robust information about aberrant brain dynamic configurations for understanding the psychopathological mechanisms in schizophrenia.

## Ethics Statement

All participants provided informed written consent. This research was approved by the respective Universities/Hospitals depending on the origin of the dataset (Medical Ethics Committee of Wuxi Mental Health Center, Nanjing Medical University for the WuXi cohort, Institutional Review Boards at UCLA and the Los Angeles County for the UCLA cohort, and institutional review board protocols of the University of New Mexico for the COBRE cohort). This study was conducted in accordance with the Declaration of Helsinki guidelines.

## Data and Code Availability Statements

The two open cohorts were obtained from UCLA Consortium for Neuropsychiatric Phenomics LA5c Study (https://openneuro.org/datasets/ds000030/versions/1.0.0) and The Center for Biomedical Research Excellence (COBRE) (http://fcon_1000.projects.nitrc.org/indi/retro/cobre.html). The WuXi cohort is not publicly available due to privacy or ethical restrictions. The code that supports the findings of this study will be made available upon request from the corresponding author.

## Acknowledgements

We thank Robert Bilder, Russell Poldrack and their colleagues for sharing the data (OpenNEURO). We are grateful to all the patients and volunteers of this study as well as the staffs at the Wuxi Mental Health Center for their help with participant recruitment and data collection. This work was supported by the National Natural Science Foundation of China (NSFC) grant (No. 62071109 to C.M., No. 81871081 and 81301148 to L.T., No. 61871420 to B.B.,).

