## supplementary materials for "Reproducible Coactivation Patterns of Functional Brain Networks Reveal the Aberrant Dynamic State Transition in Schizophrenia"

**Methods and Results**

**Subjects**

**Verification cohort 1 (COBRE):** Data used in this study was drawn from a Mind Research Network Center of Biomedical Research Excellence (COBRE), funded by the National Institutes of Health. After excluding subjects with large head motion, 54 SZ subjects and 72 HC subjects remained for the current study. A structured clinical interview based on DSM-IV was used by trained clinical psychiatrists to diagnose patients with schizophrenia in this dataset. Psychopathological symptoms were rated using the Positive and Negative Syndrome Scale (PANSS).

**Verification cohort 2 (UCLA):** Data used in this study was obtained from the OpenfMRI database with the accession number of ds000030. Detailed information about the inclusion and exclusion criteria can be found in Poldrack et al. (Poldrack et al., 2016). After excluding subjects with large head motion, 45 SZ subjects and 108 HC subjects remained for the current study. The general symptom severity was measured using the Brief Psychiatric Rating Scale (BPRS) for each schizophrenia patient. Particularly, SZ patients' negative and positive symptom severity were measured using the Scale for the Assessment of Negative Symptoms (SANS) and the Scale for the Assessment of Positive Symptoms (SAPS).

**fMRI Data Acquisition**

**Verification cohort 1 (COBRE):** A 3T Siemens Trio scanner was used to acquire rest fMRI and T1-weighted structural MRI images. The resting fMRI data were collected with single-shot full k-space echo-planar imaging (EPI) with ramp sampling correction using the intercomissural line (AC-PC) as a reference, and the parameters were followed: TR = 2000 ms, TE = 29 ms, slice number = 33, slice thickness = 3.5 mm, flip angle = 75°, matrix size = 64 × 64, FOV = 240 mm, voxel size = 3.75 × 3.75 × 4.55 mm^3^, and volume number = 150. The parameters for the T1-weighted MPRAGE structural image were the following: TR = 2.53 s, TE = 1.64 ms, slice number = 192, slice thickness = 1 mm, matrix size = 256 × 256, FOV = 256 mm, and voxel size = 1 × 1 × 1 mm^3^.

**Verification cohort 2 (UCLA):** Two 3T Siemens Trio scanners were used to acquire resting-state fMRI and T1-weighted structural MRI images. A T2*-weighted echoplanar imaging (EPI) sequence was used to collect the fMRI data, and the parameters were followed: TR = 2000ms, TE = 30 ms, slice number = 34, slice thickness = 4 mm, flip angle = 90°, matrix size = 64 × 64, FOV = 192 mm, voxel size = 3 × 3 × 3 mm^3^, and volume number = 152. The parameters for the T1-weighted MPRAGE structural image were the following: TR = 1.9 s, TE = 2.26 ms, slice number = 176, slice thickness = 1 mm, matrix size = 256 × 256, FOV = 250 mm, and voxel size = 1 × 1 × 1 mm^3^.

**The selection of clustering number**

The coactivation patterns analysis is based on temporal clustering. In this study, the clustering number was tested from k = 2 to 21, and the silhouette values were calculated for each k value. As shown in the supplementary Figure S1, it was found that the silhouette score was found to be monotonically decreasing with the increase of k value, and 2 clusters got the largest silhouette score. Then elbow criterion was considered for determining the number of clusters. While one issue is that the time points of the three datasets are limited, if the cluster number is too large, then each brain state would only account for a few seconds through the entire scan. After comprehensively considering the clustering silhouette curve for the three datasets, and the trade-off between clustering numbers and time points within each cluster, 6 clusters were mainly analyzed and reported in the manuscript.

**The relationship between different dynamic measures**

To evaluate the dynamic properties within and between CAP states, eight dynamic measures (CAP matrices) were calculated at the individual level:

(1) **Fraction of time** is defined as the proportion of total volumes spent in one CAP state over the whole time series;

(2) **Persistence** is the average time spent in one state before transferring to another state, and it describes the mean volume-to-volume maintenance of one CAP state;

(3) **Number of states (Counts)** is how many times one state occurred during the whole time series;

(4) **Transitions matrix** records how many times that State A transfers to State B, and it focuses on the macroscopic state-to-state transitions with the diagonal set to zero;

(5) **Transition probability matrix** is the probability that one volume within State A transfers to the next volume belonging to State B, with a non-zero diagonal as the volume within State A could still stay within State A for the next volume;

(6) **Resilience** is the probability or likelihood of remaining in the same state from one volume to the next volume, and it can be extracted from the diagonal of the transition probability matrix;

(7) **In-degree** and (8) **Out-degree** are the total frequencies of transition from other states to State A or from State A into other states.

However, not all of these CAP matrices are independent, and there is no necessity to analyze all the eight matrices in the same study. For instance, both persistence and resilience describe how long one state will persist before transferring to another state. We measured the relationship between the fraction of time, persistence, counts, resilience, in-degree and out-degree using the Pearson correlation. As described in the supplementary materials (Fig.S2), persistence and resilience were highly correlated; and the correlation coefficients between counts, in-degree/out-degree were almost to 1. Therefore, after excluding some dependent dynamics measures, we mainly analyzed the fraction of time, persistence, counts and transition probability in this study.

As mentioned in the manuscript, we calculated eight matrixes to measure brain state dynamics. However, these matrixes are not independent, for example, fraction of time = time spent in that state / total time, and time spent in that state = persistence * counts, **fraction of time = (persistence * counts) / total time**, here total time is a constant number. Therefore, the fraction of time is determined by both the persistence and counts.

The resilience is the probability for one volume to stay in the same state for the next volume, and the higher probability that one state would stay within itself, the longer this state would persist, hence the **persistence ∝ resilience**.

At last, the counts record the occurrences for one state. And if one state occurred during the middle of the whole time series, which means it was transferred from the previous state and would change to the next state, and this is the definition of in-degree and out-degree. Unless the occurrence of that state is at the beginning or the end, then the in-degree and out-degree would decrease one time, which equals **counts = in-degree ± 1, counts = out-degree ± 1, and in-degree = out-degree ± 1**.

The relationships between these dynamic measures were further verified using Pearson correlation, as shown in Figure S2. It can be observed that the persistence and resilience were highly correlated; The correlation coefficients between counts, in-degree and out-degree were almost to 1. However, the correlation coefficients between persistence/resilience and count/in-degree/out-degree were near to zero, which indicates how long one state will persist is independent of how many times it would occur. The fraction of time was related to persistence and counts. These results were consistent with our assumptions, therefore, we only reported the fraction of time, persistence and counts in the manuscript.

**Reproducibility and repeatability analysis**

In this study, the reproducibility and repeatability were evaluated from four aspects, three different preprocessing pipelines, four different number of ROIs, different number of clusters and three independent datasets. The results reported in the manuscript were based on the WuXi dataset, rest preprocessing with global signal regression (GSR), 408 ROIs and k = 6. This was used as the default configuration, then each time only one factor would change and the others remained the same for the new configurations.

At first, we compared the spatial similarity between states using Pearson correlation coefficients. Then, considering the label for each state would change in different configurations, the label for each state in the new configuration was changed to the corresponding label in the default configuration according to the highest spatial similarity, which could make the comparison between different configurations more straightforward. Finally, the spatial coactivation patterns maps and their dynamics were compared across different configurations. Here, we only presented the fraction of time as an example.

These results were shown in Figure S3 to Figure S10. In summary, different preprocessing pipelines and ROI numbers obtained consistent results, not only for the spatial coactivation patterns but also for the temporal dynamics group differences. With the increase of clustering number, such as from k = 6 to k = 8, two states were spatially divided, and the other states remained the same spatial coactivation patterns and temporal dynamics.

The repeatability across different datasets was lower than other configurations. As mentioned in the manuscript (Figure 6), if we applied the clustering results obtained from WuXi into the other two datasets, they showed consistent group differences in WuXi and COBRE. While the UCLA showed a different trend for the group differences, although none of them was significant after FDR correction. Nevertheless, if we compared the results obtained by the UCLA with WuXi independently, as shown in Figure S11 to Figure S12, they showed considerable spatial overlaps. For instance, state 1 and state 2 in WuXi were similar to state 2 and state 3 in UCLA, all of them were dominated by the frontal-parietal network, furthermore, their showed consistent group differences in fraction of time. Therefore, there was still considerable repeatability across different datasets.

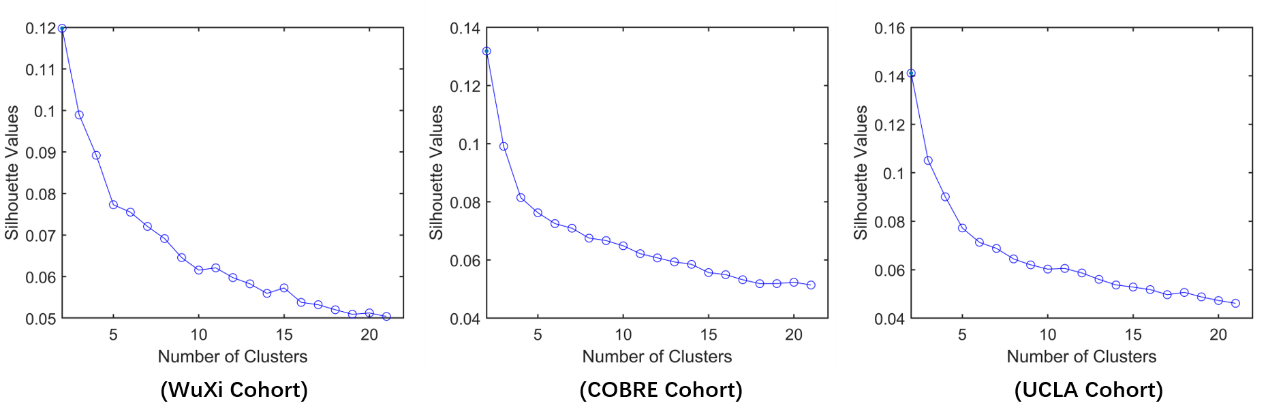

**Figure S1.** The clustering curve in the three datasets. Silhouette value was calculated from k = 2 to k =21 with step length = 1, and the silhouette values were monotonically decreasing with the increase of k. The elbow point for the three curves was around 4 to 6, and k = 6 was chosen in the manuscript.

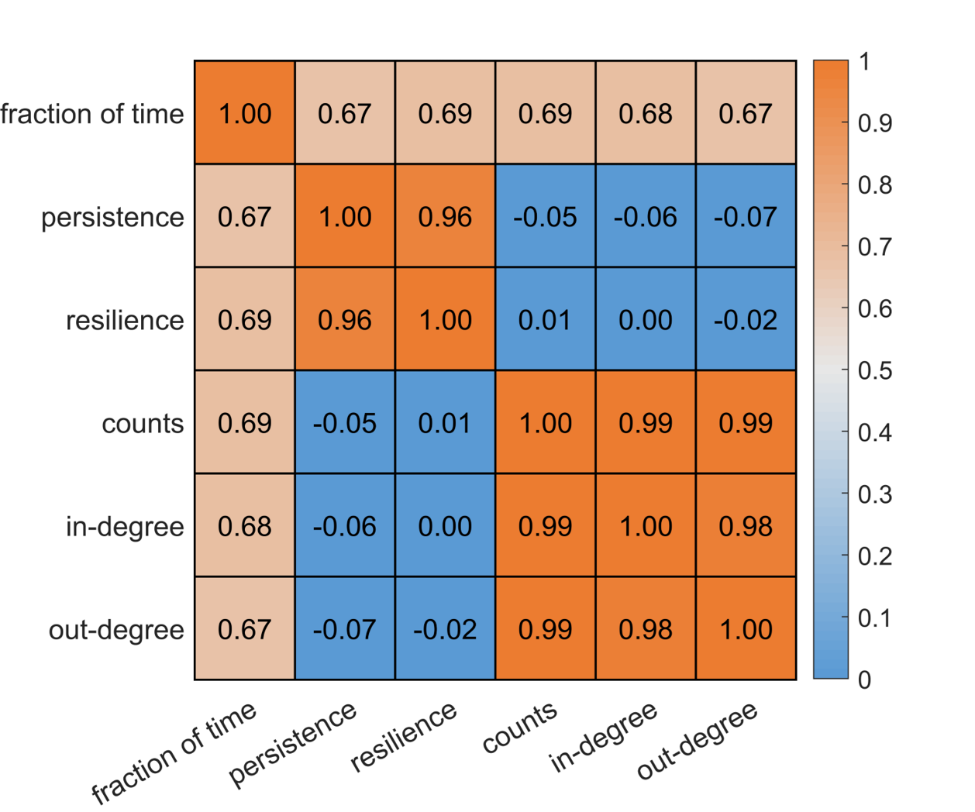

**Figure S2.** The relationship between different dynamics matrices. It can be observed that the persistence and resilience were highly correlated; The correlation coefficients between counts, in-degree and out-degree were almost to 1. However, the correlation coefficients between persistence/resilience and count/in-degree/out-degree were near to zero, which indicates how long one state will persist is independent of how many times it would occur. The fraction of time was related to persistence and counts.

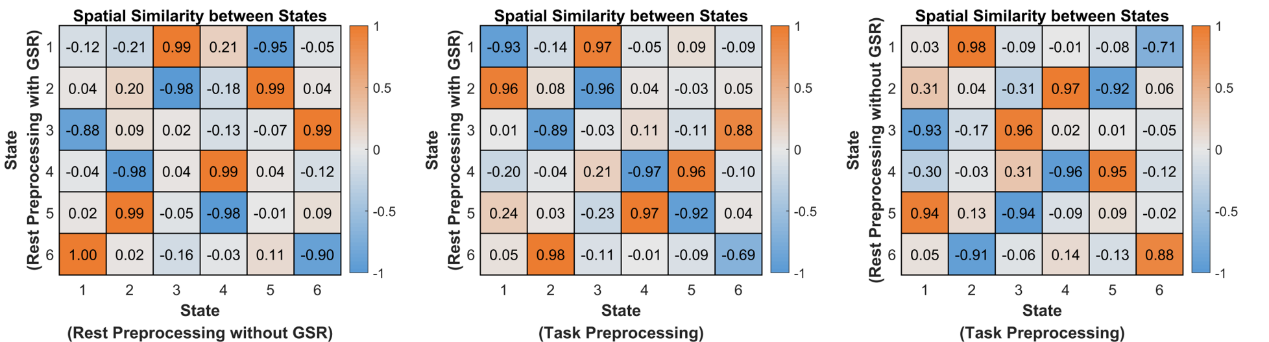

**Figure S3.** The spatial similarity between states for different preprocessing pipelines. In this case, the configuration was WuXi cohort, 408 ROIs were used, and k = 6. Different preprocessing pipelines achieved a good spatial consistency, and each state in the new configuration has a corresponding unique state in the default configuration.

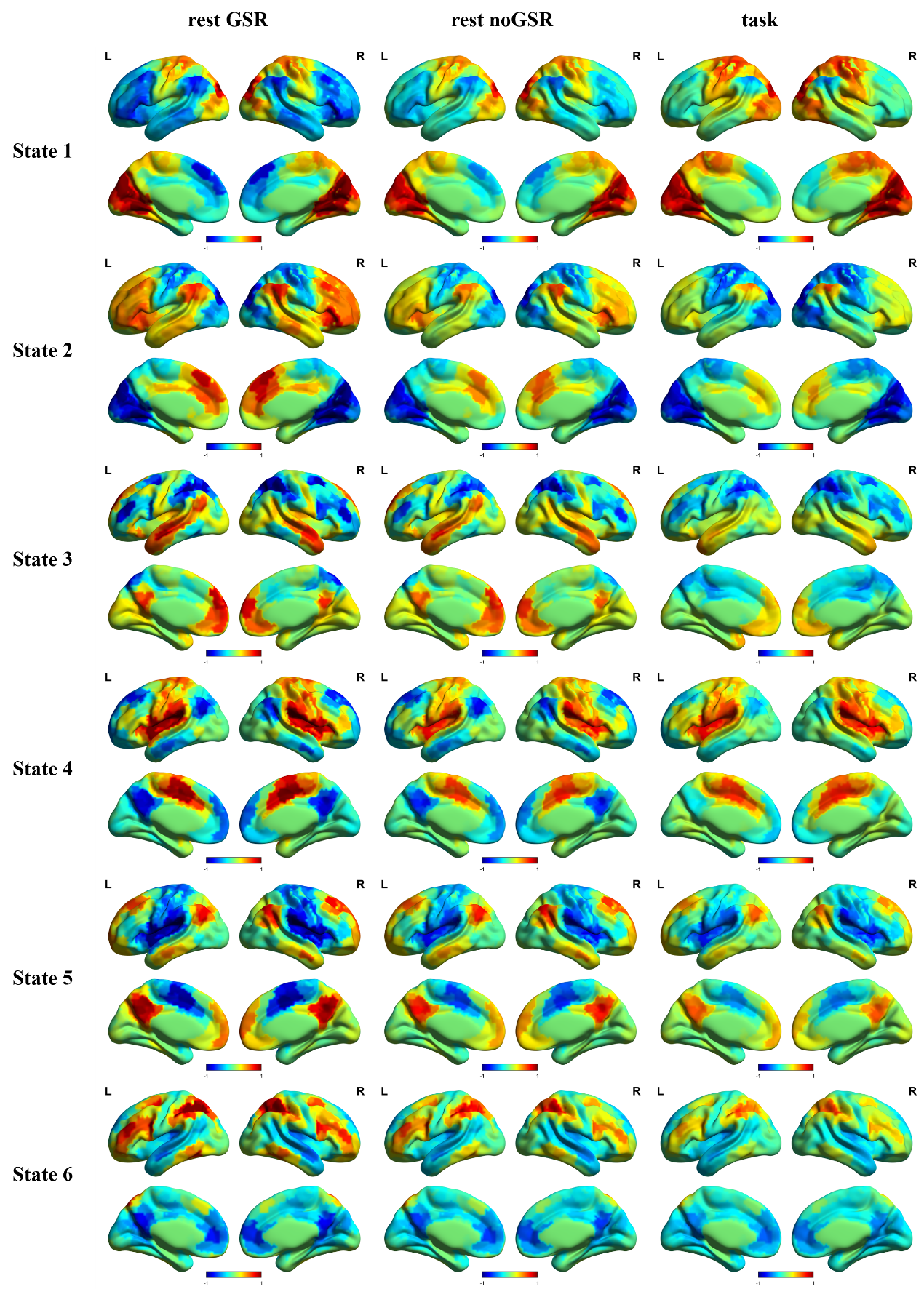

**Figure S4.** The coactivation patterns for different preprocessing pipelines. In this case, the configuration was WuXi cohort, 408 ROIs were used, and k = 6. The first and the second columns were based on the rest preprocessing pipeline with and without global signal regression (GSR), and the third column was based on the task preprocessing pipeline. For all the six states, the spatial coactivation patterns were similar across different preprocessing pipelines, and the rest preprocessing pipeline with GSR has more contrast between positive and negative activations.

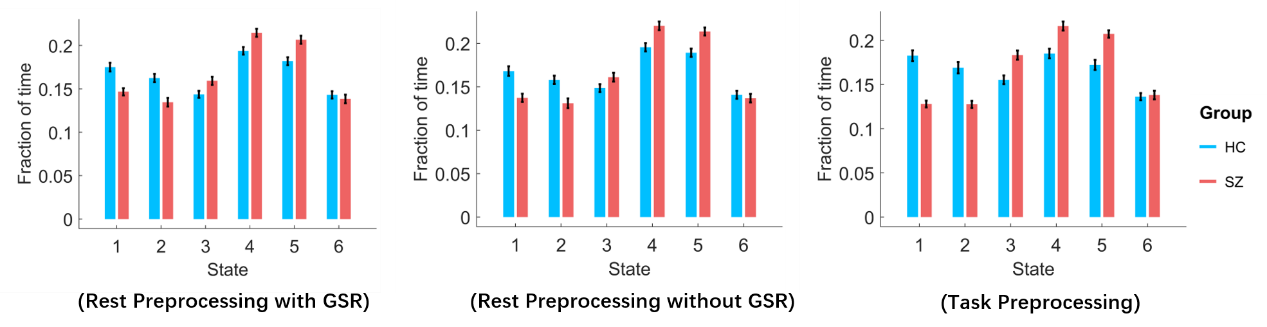

**Figure S5.** The fraction of time for different preprocessing pipelines. In this case, the configuration was WuXi cohort, 408 ROIs were used and k = 6. The first and the second columns were based on the rest preprocessing pipeline with and without global signal regression (GSR), and the third column was based on the task preprocessing pipeline. The trend for the group differences was similar across different preprocessing pipelines. For example, SZ showed less fraction of time in state 1 and state 2, and more fraction of time in state 4 and state 5.

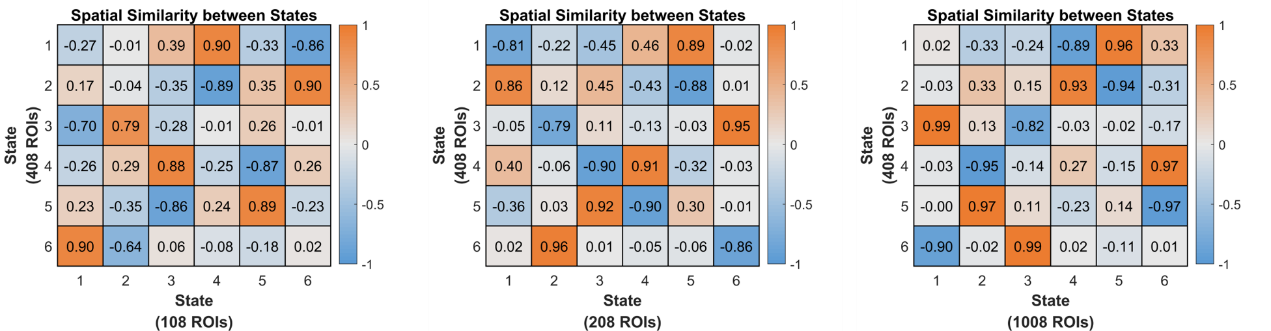

**Figure S6.** The spatial similarity between states for different number of ROIs. In this case, the configuration was WuXi cohort, rest preprocessing with GSR and k = 6. Different number of ROIs achieved a good spatial consistency, and each state in the new configuration has a corresponding unique state in the default configuration.

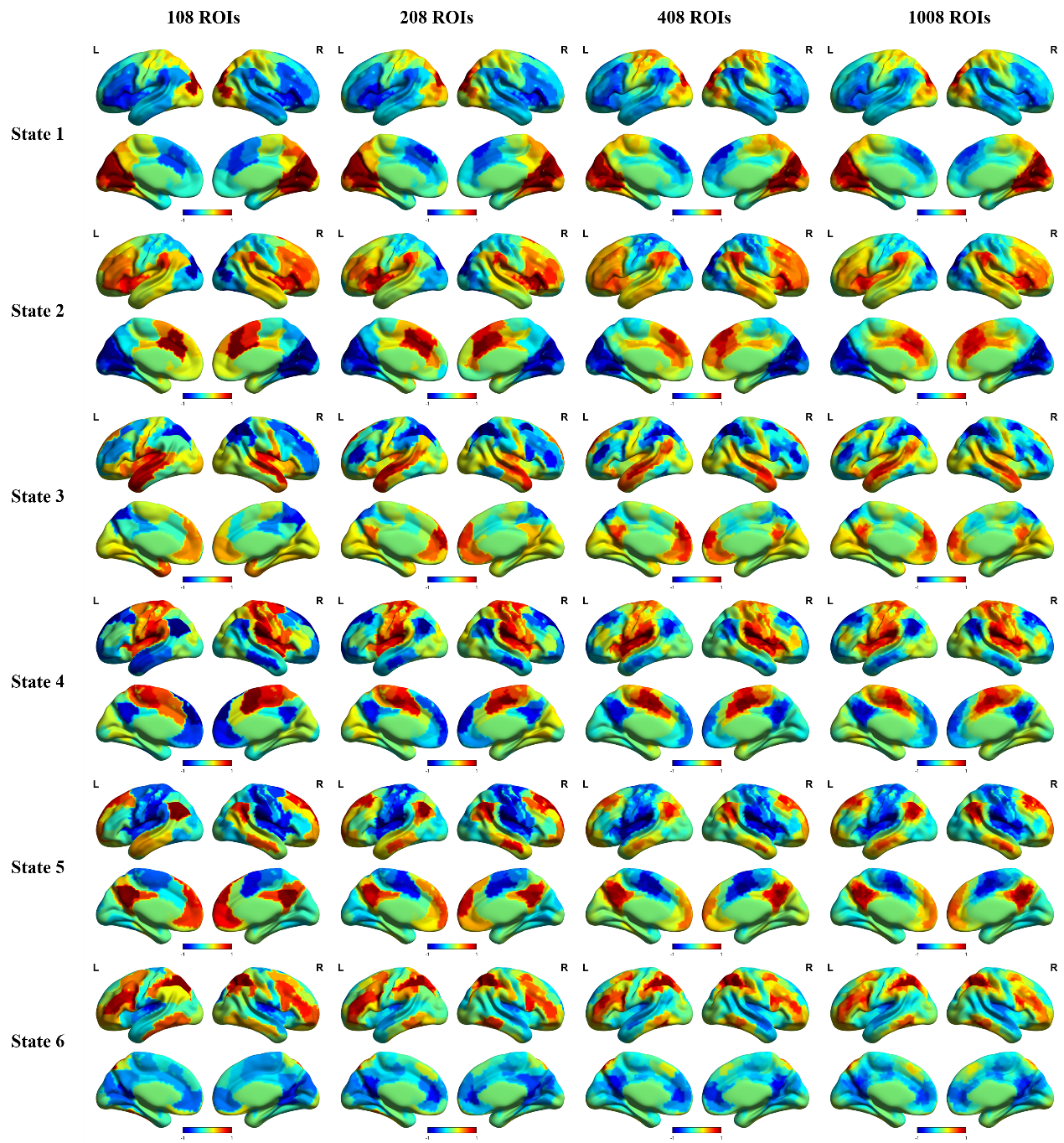

**Figure S7.** The coactivation patterns for different number of ROIs. In this case, the configuration was WuXi cohort, rest preprocessing with GSR and k = 6. The four columns were based on 108, 208, 408 and 1008 ROIs. For all the six states, the spatial coactivation patterns were similar across different number of ROIs, and the 108 ROIs has more contrast between positive and negative activations.

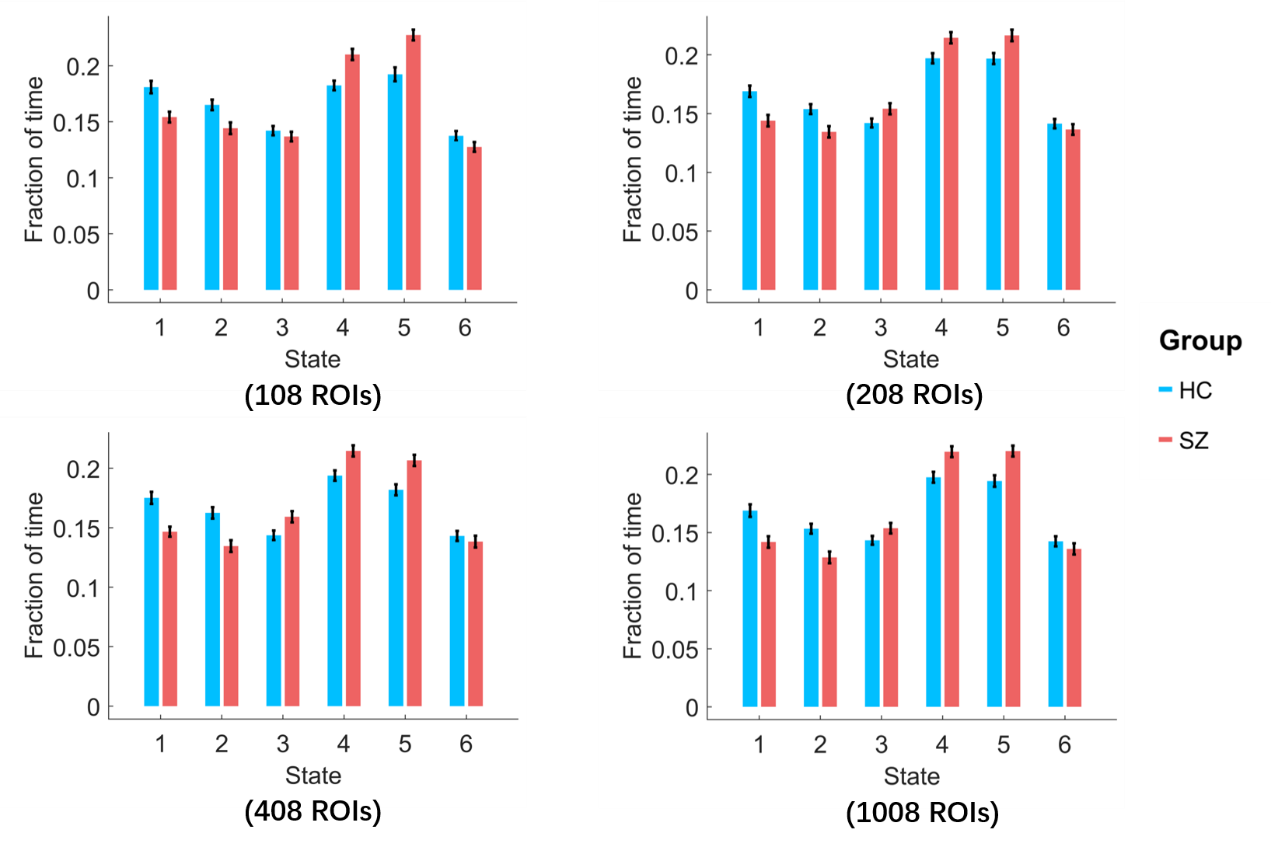

**Figure S8.** The fraction of time for different number of ROIs. In this case, the configuration was WuXi cohort, rest preprocessing with GSR and k = 6. The trend for the group differences was similar across different number of ROIs. For example, SZ showed less fraction of time in state 1 and state 2, and more fraction of time in state 4 and state 5.

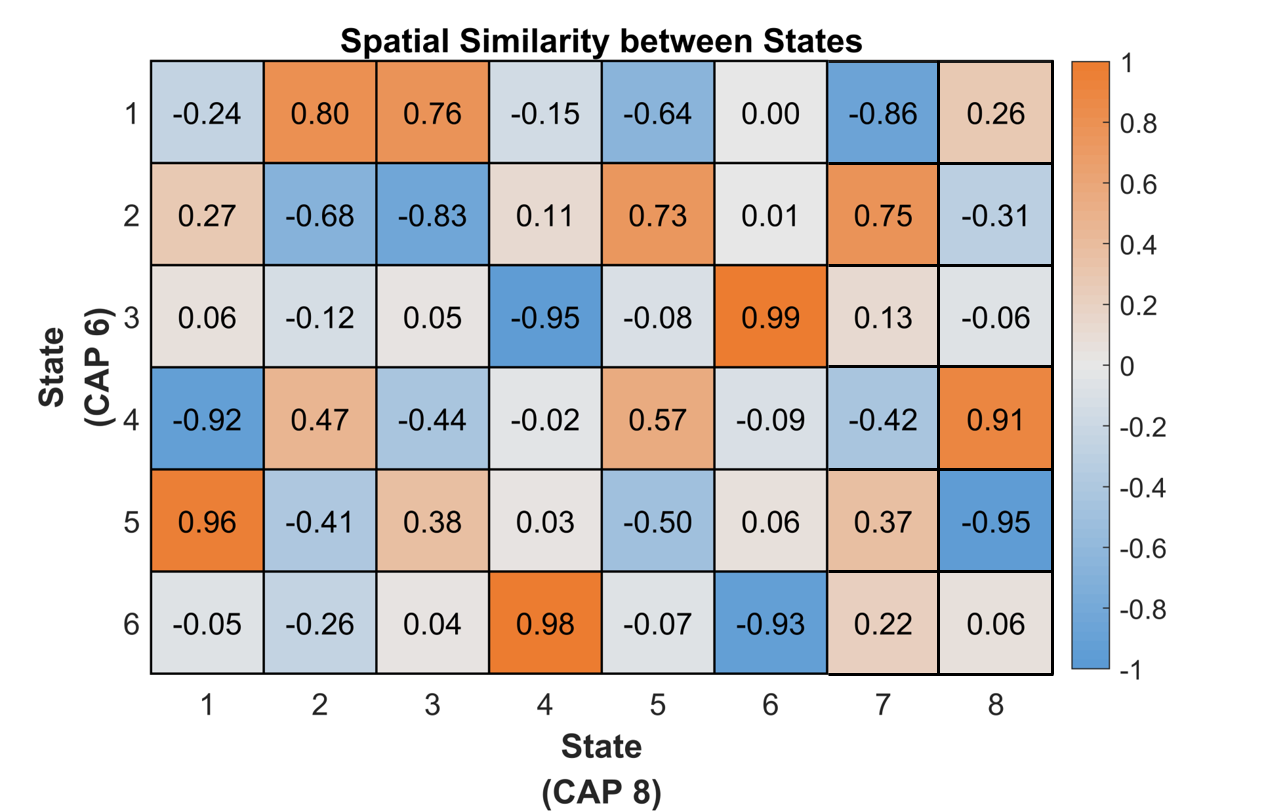

**Figure S9.** The spatial similarity between states for different numbers of clusters. In this case, the configuration was WuXi cohort, rest preprocessing with GSR, and 408 ROIs. The column is k = 6 and the row is k = 8. It can be observed that for k = 6, state 3 to state 6 has one unique corresponding state in k = 8, while state 1 (k = 6) was divided into state 2 and state 3 (k = 8), and state 2 (k = 6) was divided into state 5 and state 6 (k = 8).

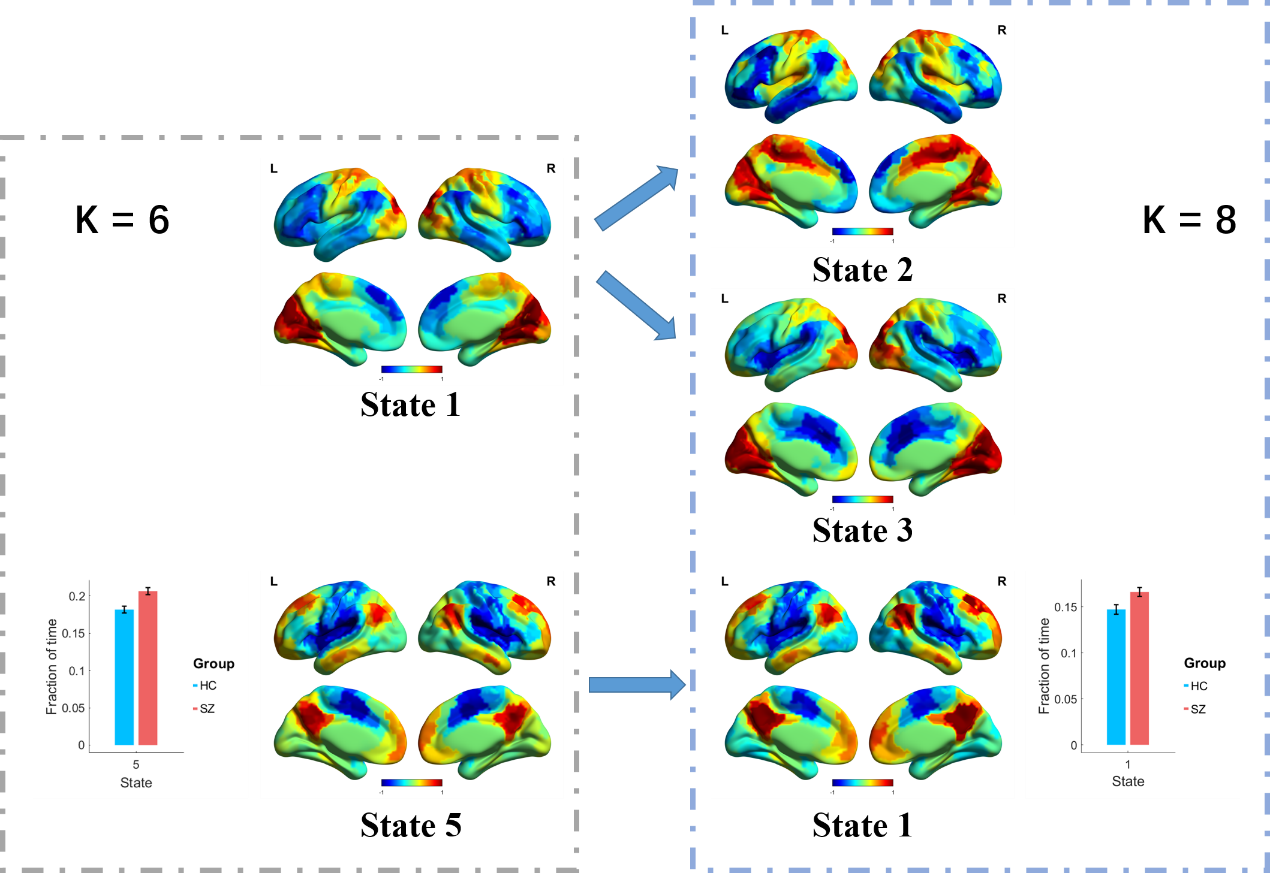

**Figure S10.** The coactivation patterns and the fraction of time for k = 6 and k = 8. In this case, the configuration was WuXi cohort, rest preprocessing with GSR and 408 ROIs. As shown in Figure S9, state 1 in k = 6 was divided into two states, state 2 and state 3 in k = 8. The state 5 in k = 6 remained in k = 8, which corresponds to state 1, and the SZ group showed a consistent more fraction of time than the HC group.

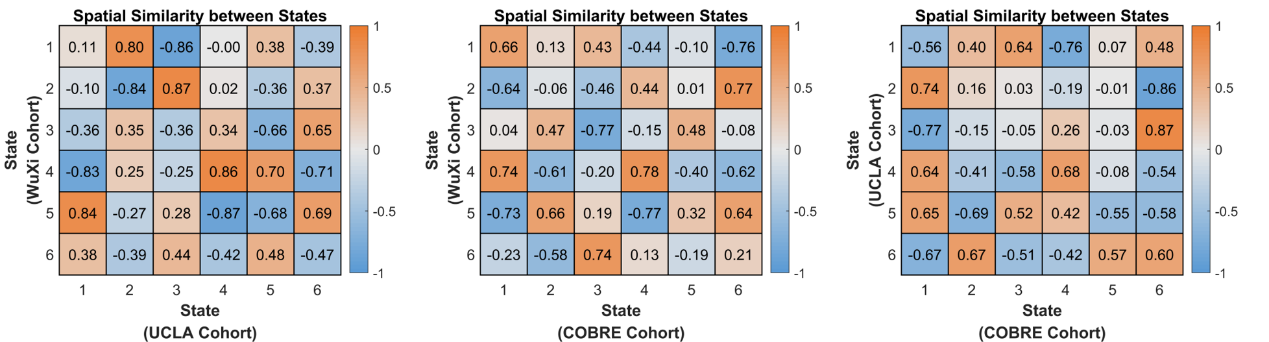

**Figure S11.** The spatial similarity between states for different cohorts. In this case, the configuration was rest preprocessing with GSR, 408 ROIs and k = 6. Compared with previous results which changed the preprocessing pipeline, number of cluster or k, the three independent datasets showed less spatial similarity between states, and not all the six states have the one-one correspondence between datasets.

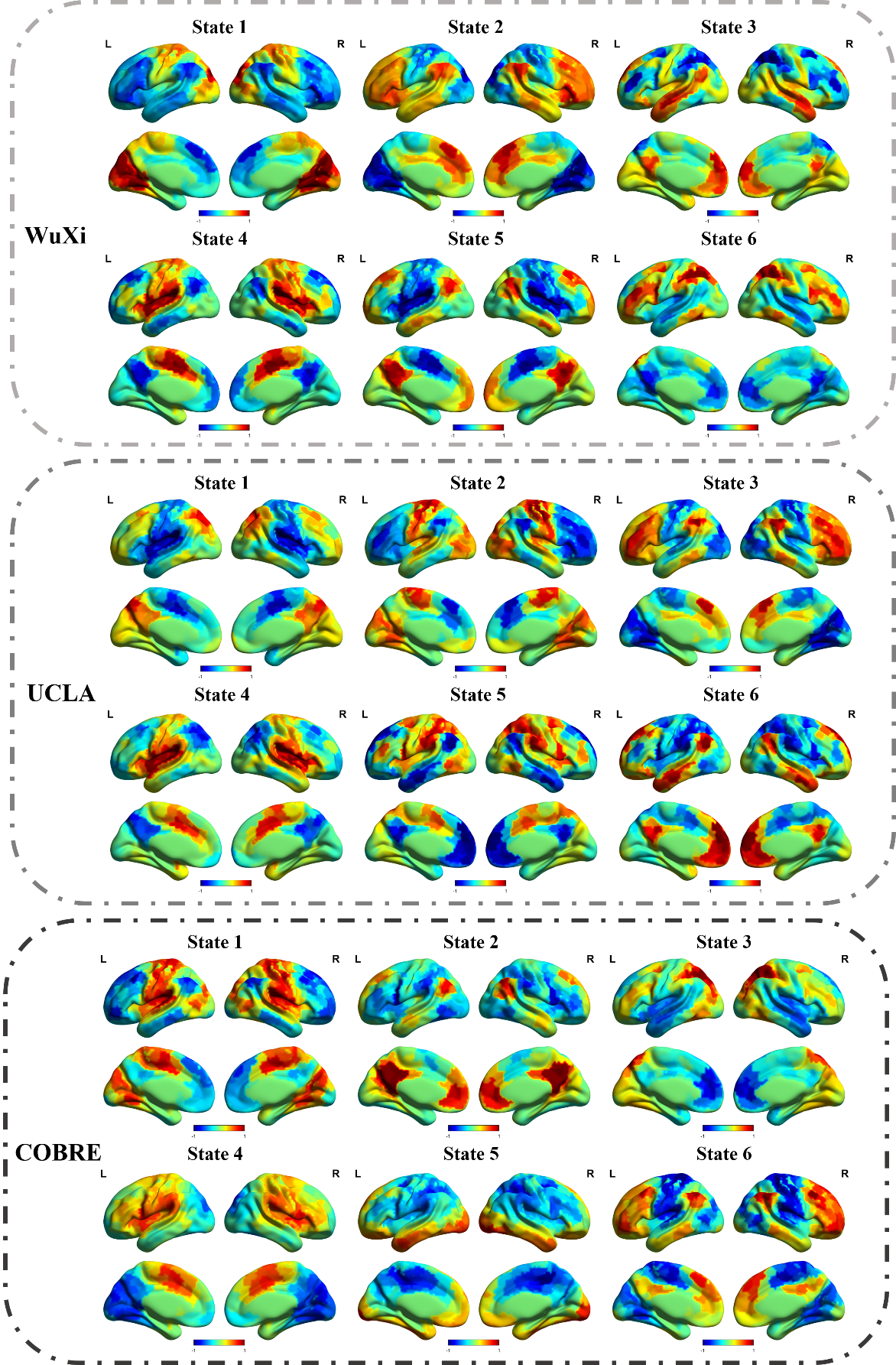

**Figure S12.** The coactivation patterns for different cohorts. In this case, the configuration was rest preprocessing with GSR, 408 ROIs and k = 6.

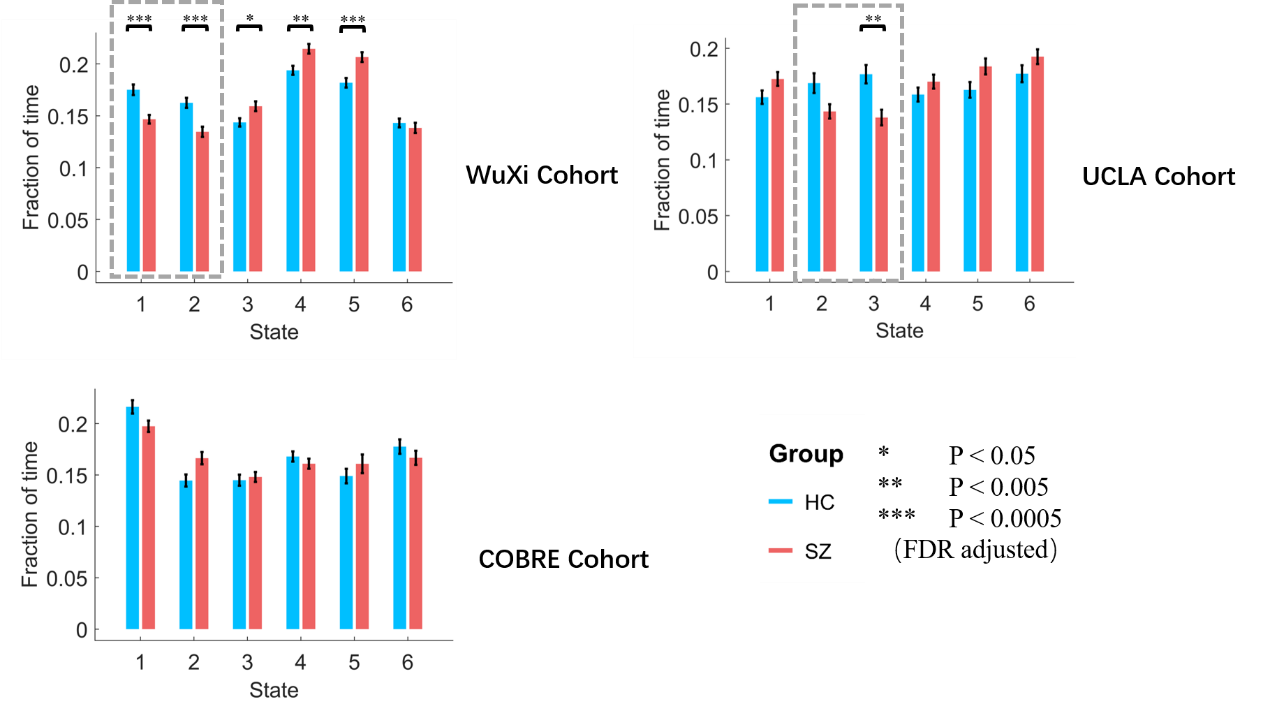

**Figure S13.** The fraction of time for different cohorts. In this case, the configuration was rest preprocessing with GSR, 408 ROIs and k = 6. * indicates p < 0.05, ** indicates p < 0.005 and *** indicates p < 0.005 (FDR adjusted). For instance, the state 1 and state 2 in WuXi were similar to state 2 and state 3 in UCLA, all of them were dominated by the frontal-parietal network, furthermore, they showed consistent group differences in fraction of time.

**Table S1.** The full demographic information for the three datasets

| Dataset | Preprocessed | Statistics | | |
| --- | --- | --- | --- | --- |
| WuXi | HC  (n = 97)  Mean ± SD | HC  (n = 69)  Mean ± SD | SZ  (n = 69)  Mean ± SD | P value |
| Age | 40.36 ± 14.77 | 45.84 ± 11.89 | 46.06 ± 10.96 | 0.9112 |
| Gender (M \ F) | 56 \ 41 | 35 \ 34 | 35 \ 34 | 1 |
| Disease duration | - | - | 19.84 ± 10.96 | - |
| PANSS positive | - | - | 20.06 ± 4.59 | - |
| PANSS negative | - | - | 23.78 ± 3.84 | - |
| COBRE | HC  (n = 72)  Mean ± SD | HC  (n = 54)  Mean ± SD | SZ  (n = 54)  Mean ± SD | P value |
| Age | 35.85 ± 11.54 | 37.22 ± 12.48 | 37.80 ± 14.13 | 0.8234 |
| Gender (M \ F) | 49 \ 23 | 43 \ 11 | 43 \ 11 | 1 |
| Disease duration | - | - | 15.19 ± 12.46 | - |
| PANSS positive | - | - | 14.13 ± 4.29 | - |
| PANSS negative | - | - | 14.46 ± 5.10 | - |
| UCLA | HC  (n = 108)  Mean ± SD | HC  (n = 45)  Mean ± SD | SZ  (n = 45)  Mean ± SD | P value |
| Age | 31.20 ± 8.77 | 36.73 ± 8.65 | 37.00 ± 8.75 | 0.8973 |
| Gender (M \ F) | 60 \ 48 | 33 \ 12 | 33 \ 12 | 1 |
| BPRS | - | - | 50.40 ± 14.08 | - |
| SAPS | - | - | 28.89 ± 18.49 | - |
| SANS | - | - | 35.09 ± 18.55 | - |

Preprocessed: All HC subjects remained after quality control (e.g., head motion), and all HC subjects were used in the k-means clustering analysis to get the coactivation patterns.

Statistics: Only the age- and gender-matched HC subjects were selected to compare with SZ patients.

Abbreviations: BPRS, Brief Psychiatric Rating Scale; SANS, Scale for the Assessment of Negative Symptoms; SAPS, Scale for the Assessment of Positive Symptoms; SD: Standard Deviation; PANSS, Positive and Negative Syndrome Scale;

Statistics: Age, two-sample t-test; Gender, chi-square cross-table test.

**Table S2.** The two-sample t-test for network dynamics measures differences between SZ and HC. The configuration was WuXi cohort, rest preprocessing with GSR, 408 ROIs and K = 6.

| Fraction of time | T value | P value (FDR adjusted) |
| --- | --- | --- |
| State 1 | -4.3513 | 0.0002 |
| State 2 | -4.0603 | 0.0002 |
| State 3 | 2.5183 | 0.0156 |
| State 4 | 3.2579 | 0.0021 |
| State 5 | 3.8056 | 0.0004 |
| State 6 | -0.7270 | 0.4685 |
| Persistence | T value | P value (FDR adjusted) |
| State 1 | -4.0973 | 0.0004 |
| State 2 | -2.6912 | 0.0241 |
| State 3 | -0.4947 | 0.6216 |
| State 4 | 0.9752 | 0.3975 |
| State 5 | 1.9984 | 0.0954 |
| State 6 | -1.0484 | 0.3975 |
| Counts | T value | P value (FDR adjusted) |
| State 1 | -0.8557 | 0.4724 |
| State 2 | -3.1975 | 0.0052 |
| State 3 | 3.3235 | 0.0052 |
| State 4 | 2.8751 | 0.0094 |
| State 5 | 2.5688 | 0.0169 |
| State 6 | 0.0004 | 0.9997 |
| Transition probability | T value | P value (FDR adjusted) |
| State 1 to State 1 | -4.4354 | 0.0001 |
| State 2 to State 2 | -2.6660 | 0.0259 |
| State 1 to State 5 | 2.8039 | 0.0452 |
| State 2 to State 3 | 2.7132 | 0.0452 |
| State 3 to State 5 | 2.7198 | 0.0452 |
| State 4 to State 2 | -2.9329 | 0.0452 |
| State 6 to State 4 | 3.0510 | 0.0452 |

Notes: the within state transition probability (resilience) was corrected separately.
